# Subtractive proteomics unravel the potency of D-Alanine-D-Alanine Ligase as the drug target for *Burkholderia pseudomallei*

**DOI:** 10.1101/2024.10.15.618487

**Authors:** Shakilur Rahman, Ayush Bhattacharya, Prerona Jana, Manisha Ganguly, Amit Kumar Das, Ditipriya Hazra, Amlan Roychowdhury

## Abstract

Melioidosis, also known as Whitmore’s disease, is caused by the deadly pathogen *Burkholderia pseudomallei* and remains a significant global health concern, particularly in South Asia. The disease is contracted through exposure to contaminated soil, water, air, and food. Infected individuals often present with abscesses in internal organs such as the lungs, spleen, and liver, and in soft tissues, with severe cases leading to septic shock and acute pneumonia. The rising incidence and mortality rates, coupled with *B. pseudomallei’s* ability to form biofilms and develop resistance to antibiotics like cephalosporins, make treatment increasingly challenging. This highlights the urgent need for novel therapeutic approaches.

D-Alanine-D-Alanine ligase (Ddl), a crucial enzyme involved in the final stage of bacterial cell wall synthesis, which protects the pathogen from the hostile cellular environment of the host. While many bacteria have two isoforms of this enzyme, *B. pseudomallei* possesses only the DdlB isoform, presenting a significant vulnerability. Our study represents the first successful attempt to target DdlB through a combination of molecular docking and molecular dynamics simulations. These investigations provide strong evidence that Conivaptan acts as an effective inhibitor of DdlB, offering a novel therapeutic approach for combating melioidosis.

## INTRODUCTION

Melioidosis, caused by the bacterium *Burkholderia pseudomallei*, is a serious global health threat first identified by Alfred Whitmore in Myanmar over a century ago (Whitmore, 1913). Commonly known as Whitmore’s Disease, melioidosis is contracted through inhalation, skin cuts, and occasionally ingestion. The disease primarily affects adults with underlying health conditions, such as diabetes mellitus, which impairs immune responses (Wiersinga et al., 2018; Cheng & Currie, 2005). Clinically, melioidosis often presents with abscesses in internal organs such as the lungs, spleen, and liver, and in soft tissues. Severe cases can lead to septic shock, frequently associated with acute pneumonia. Asymptomatic infections can progress to acute melioidosis many years after initial exposure, with the longest documented incubation period being 62 years (Dance, 2000; Gassiep et al., 2020). In children in Southeast Asia, melioidosis commonly presents as parotid abscesses. Additionally, *B. pseudomallei* is a significant veterinary pathogen, causing lethal infections in various species and exhibiting symptoms similar to glanders (Ngauy et al., 2005; Novo et al., 2003).

Globally, melioidosis accounts for approximately 165,000 cases annually, with South Asia contributing 44% of this burden (Chewapreecha et al., 2017). The disease results in 4.64 million disability-adjusted life-years (DALYs), with nearly 99% of these as years of life lost (YLL) due to high mortality rates, especially in rural, low, and middle-income populations (Birnie et al., 2019).

Antibiotics are the primary treatment for bacterial infections, but *Burkholderia pseudomallei* exhibits intrinsic and natural drug resistance mechanisms, complicating treatment efforts. β-lactams, which inhibit cell wall biosynthesis, are commonly used, but *B. pseudomallei* expresses the β-lactamase gene penA located on chromosome 2, conferring resistance to antibiotics like ceftazidime (Thonglao et al., 2022) and clavulanate (Schweizer, 2012). Additionally, biofilm formation induces resistance and causes recurrence of melioidosis (Sawasdidoln et al., 2010; Sýkorová et al., 2020). These factors necessitate the discovery of new targets that can effectively combat the pathogen without the risk of developing resistance. Recent attempts to develop vaccines have not been successful, indicating a need for further investigation (Alshabrmi & Alatawi, 2024).

Among several bacterial proteins indispensable for survival and pathogenicity, D-Alanine-D-Alanine ligase (Ddl) has acted as a drug target in many pathogenic organisms, for instance, *Mycobacterium tuberculosis* (Bruning et al., 2011), *Aeronomus hydrophila* (Zhang et al., 2020), *Staphylococcus aureus* (Liu et al., 2006), *Acinetobacter baumanii* (Ahmad et al., 2018) etc. Ddl is an ATP-dependent enzyme, produces D-alanyl-D-alanine, which forms the terminal dipeptide of the peptidoglycan monomer unit (Bouhss et al., 2008; Bugg et al., 2011). This terminal dipeptide is crucial for transpeptidation, the cross-linking process of peptidoglycan chains. Disrupting D-alanyl-D-alanine biosynthesis weakens the bacterial cell wall, leading to cell lysis, making Ddl a valuable target for antibacterial drug discovery. Ddl exists as two isoforms, DdlA and DdlB, in some bacteria, in *E*.*coli*, EcDdlA and EcDdlB shares 30% sequence identity and exhibit similar substrate specificity and inhibitor susceptibility. However, *B. pseudomallei* has only the DdlB isoform (BpDdl), presenting a unique target for drug development(al-Bar et al., 1992).

BpDdl is an ATP depenedent ligase containing three domains: the N-terminal α/β-domain, the central domain and the C-terminal domain that make up the ATP grasp fold. The catalytically active BpDdl features an ATP-binding pocket and a D-alanine binding pocket, with two Mg^2+^ ions anchoring the ATP phosphate group. The C-terminal domain has a flexible lid loop that facilitates the enzyme’s function by undergoing significant conformational changes during catalysis. Given that *B. pseudomallei* has only the DdlB isoform, it presents a unique vulnerability. Previous attempts to target Ddl using D-boroAlanine (D-Ala with the -COOH group replaced with a -B(OH)2 group) showed potent antibacterial activity, but efficacy deteriorated upon drug purification, and resistance mechanisms emerged. Therefore, there are currently no effective inhibitors of BpDdl and at the same time it also refers that finding new drug targets and target centric search of inhibitor/lead molecules are two dire needs for finding new treatments for melioidosis. (Ameryckx et al., 2018; Díaz[Sáez et al., 2019).

To address the issues two different methodologies were adopted in this study, one is using subtractive proteomics to find best possible targets. The second is brute force drug screening, target-drug interaction analysis coupled with molecular dynamics simulation and MM-PBSA based binding free energy calculation to find effective drug/lead molecule against *B. pseudomallei*. The workflow of this study has been summarized in Figure 1.

**Figure 1:**
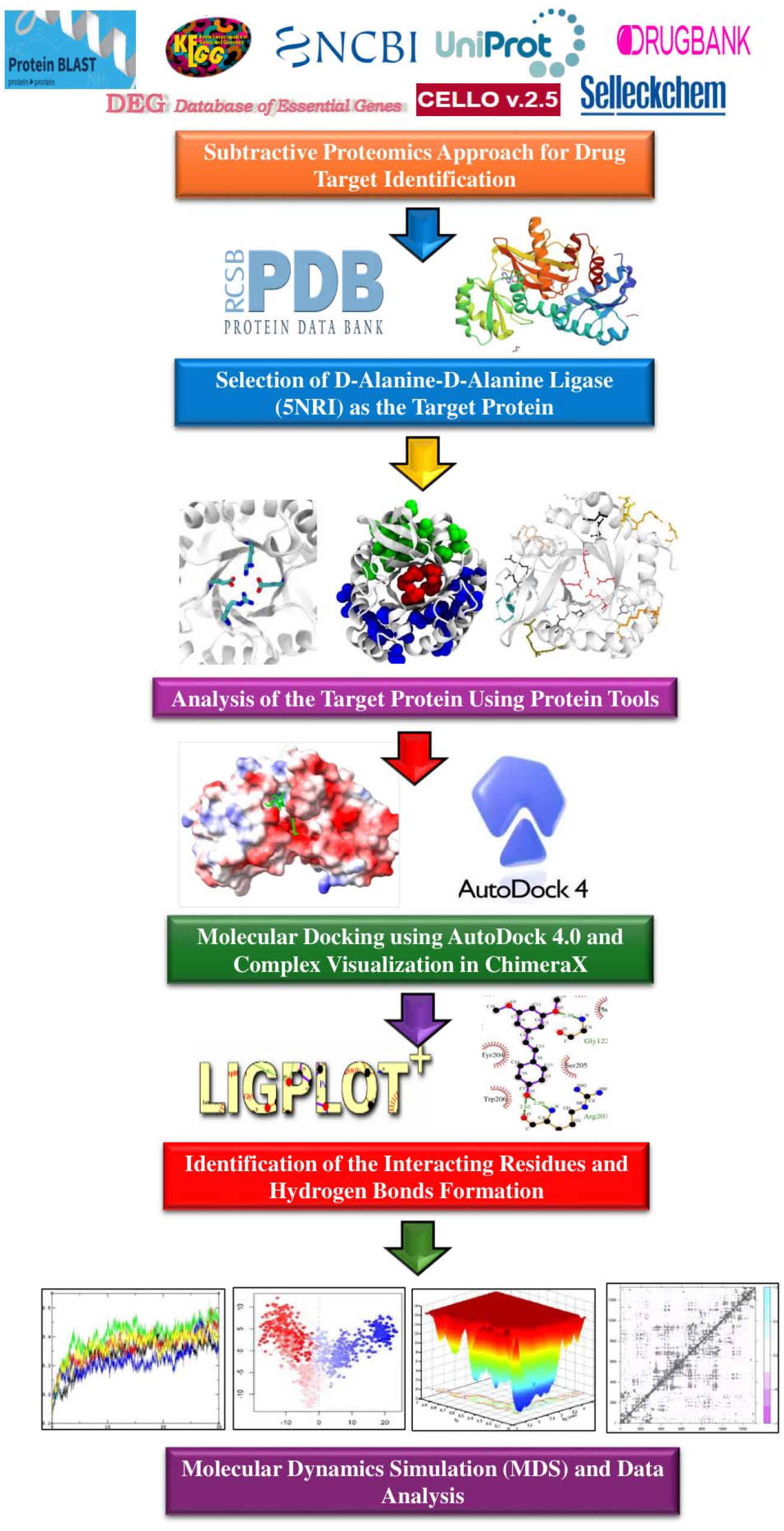
The graphical representation of the workflow of the study that represent the subtractive proteomics to investigate the possible targets against BpDdl, that was further followed by brute force screening and protein-inhibitor analysis coupled with MD simulations and binding energy calculations.

## RESULTS

### Pathogenic strain K96243 has 57 Unique Metabolic Pathways

Information about 138 metabolic pathways of *Burkholderia pseudomallei* K96243 (KEGG organism: bps) and 356 metabolic pathways of Humans (KEGG organism: *hsa*) were curated from KEGG Database. Among those, 53 unique metabolic pathways (Supplementary table 1) were observed in K96243 and 81 of the metabolic pathways (Supplementary table 2) were common between human host and pathogen (Figure 2).

**Figure 2:**
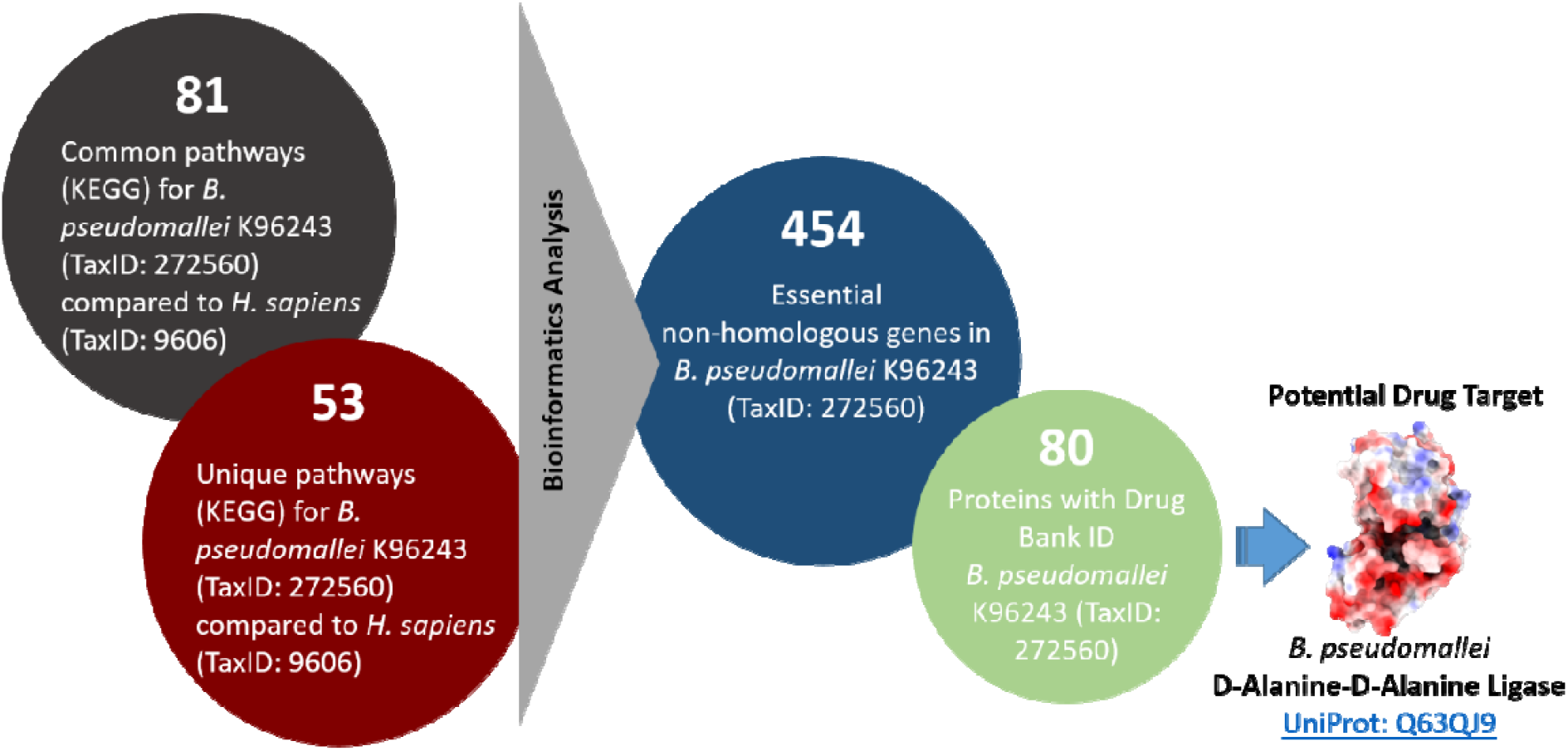
Graphical presentation of the statistics obtained from subtractive proteomics and the detail of drug target identified.

### Identification of Essential Non-Homologous Proteins

Among the 30,878 essential proteins curated from the DEG Database (as on 1st September 2020), 454 proteins were identified (Figure 2) to be essential for the survival of the bacteria and were not homologous with the human host (Supplementary table 3). These proteins were annotated as “Non-Homologous Essential Proteins”. The term itself explains the therapeutic potential of these proteins along with avoiding chances of cross reactivity in the host.

### Identification of Potential Vaccines Targets

The subcellular localization of proteins is an important factor for determining potential vaccine targets, the extracellular and secretory proteins are usually selected as vaccine candidates. In this study 25 extracellular proteins that are required for virulence or survival of the pathogen were selected as potential vaccine targets. Those 25 extracellular proteins were subjected to antigenicity prediction, to narrow down the selection for vaccine target. Antigenicity refers to the specific recognition of a substance by antibodies generated in response to the immune system. VaxiJen v2.0 is a tool utilized to assess the antigenicity of epitopes, aiding in understanding their immunomodulatory effects. Epitopes with antigenicity values exceeding the VaxiJen cut-off value of 0.4 are considered potential epitopes, typically ranging between 0.5 to 1.43 (Supplementary table 4).

### Targeting Non-Homologous Essential Proteins for Drug Repurposing

Further, the cytoplasmic non-homologous essential proteins underwent comparative genomics analysis to identify potential drug targets (Figure 2 & Supplementary table 5). A thorough literature study coupled with the analysis indicated D-Alanine-D-Alanine Ligase is a potential drug target. The structure of *Burkholderia pseudomallei* D-Alanine-D-Alanine Ligase has been portrayed in Figure 3 in cartoon mode (Figure 3A) as well as in surface (Figure 3B) where the D-Alanine binding site (potential pocket for drug screening) is marked.

**Figure 3:**
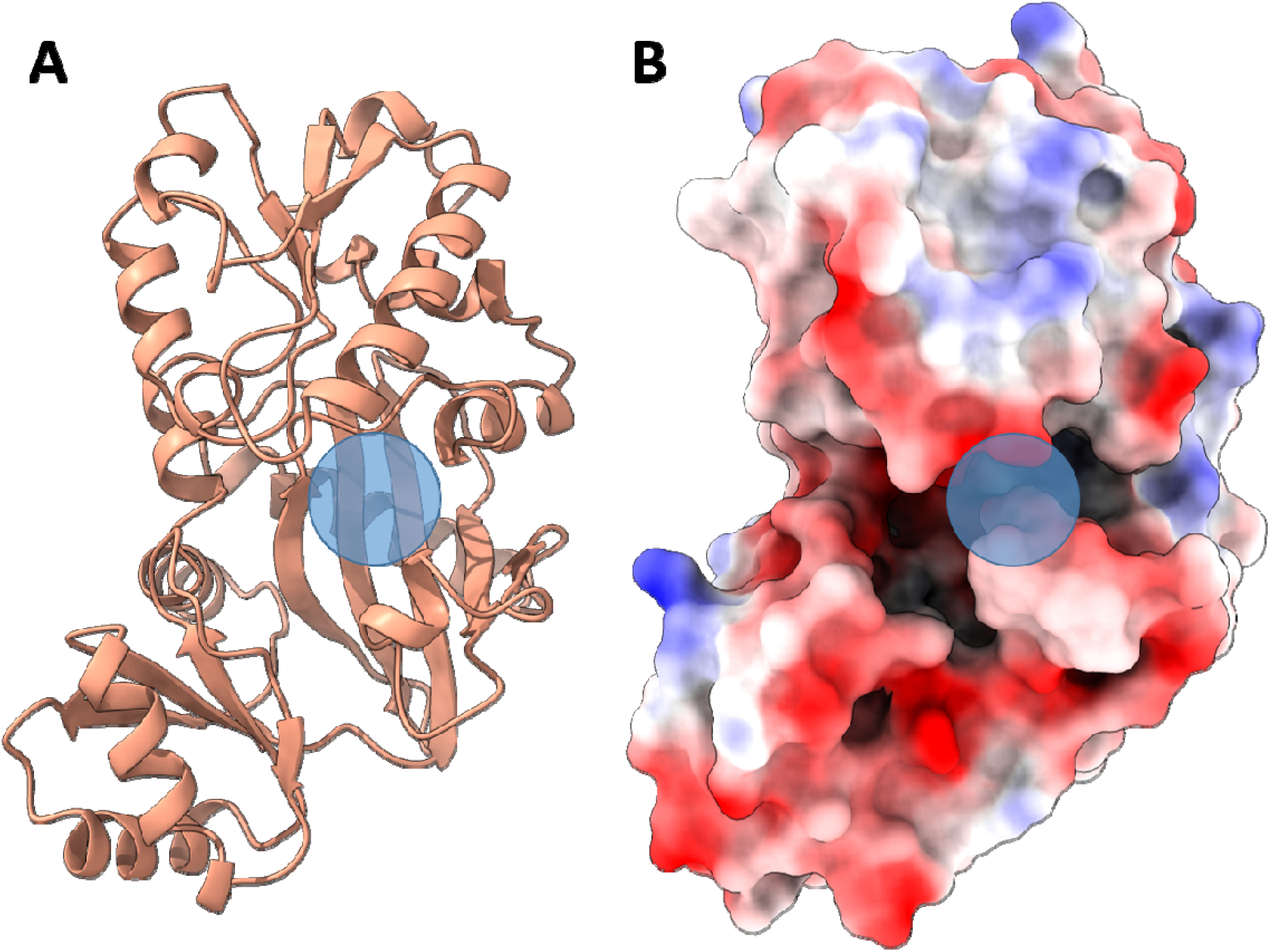
The crystal structure of BpDdl (PDB code 5NRI). (A) The cartoon representation of BpDdl structure the substrate binding site is highlighted with the semitransparent blue circle. (B) Surface representation of BpDdl structure where negatively charged regions are shown in red and the positively charged regions are in blue. The binding pocket is marked with the the semitransparent blue circle.

### Homology Modelling of D-Alanine-D-Alanine Ligase

The BpD-alanine-D-alanine Ligase crystal structure of (PDB: 5NRI) was obtained from PDB. The correct any missing residue the protein was modelled using Swiss model (Waterhouse et al., 2018).

### Identification of Interacting Partners Using StringDB

The results from the String Database (Szklarczyk et al., 2023) (Figure 4) show the bacterial protein D-Alanine-D-Alanine Ligase interacts with a variety of proteins which includes alanine racemase that take part in interconversion of L-alanine to D-alanine, cell division protein FtsA and FtsQ that plays essential role in Z ring formation and bacterial cell division respectively. The protein also interacts with various other proteins that are involved in cell wall biogenesis like mur C, mur D, mur E, mur F and mur G. The StringDB also provides information regarding the enriched pathways based on the interacting partners of the query protein. It was found that the mur family of proteins that exhibit strong interaction with BpDdl is also involved in development of Vanomycin resistance, according to the kegg data.

**Figure 4:**
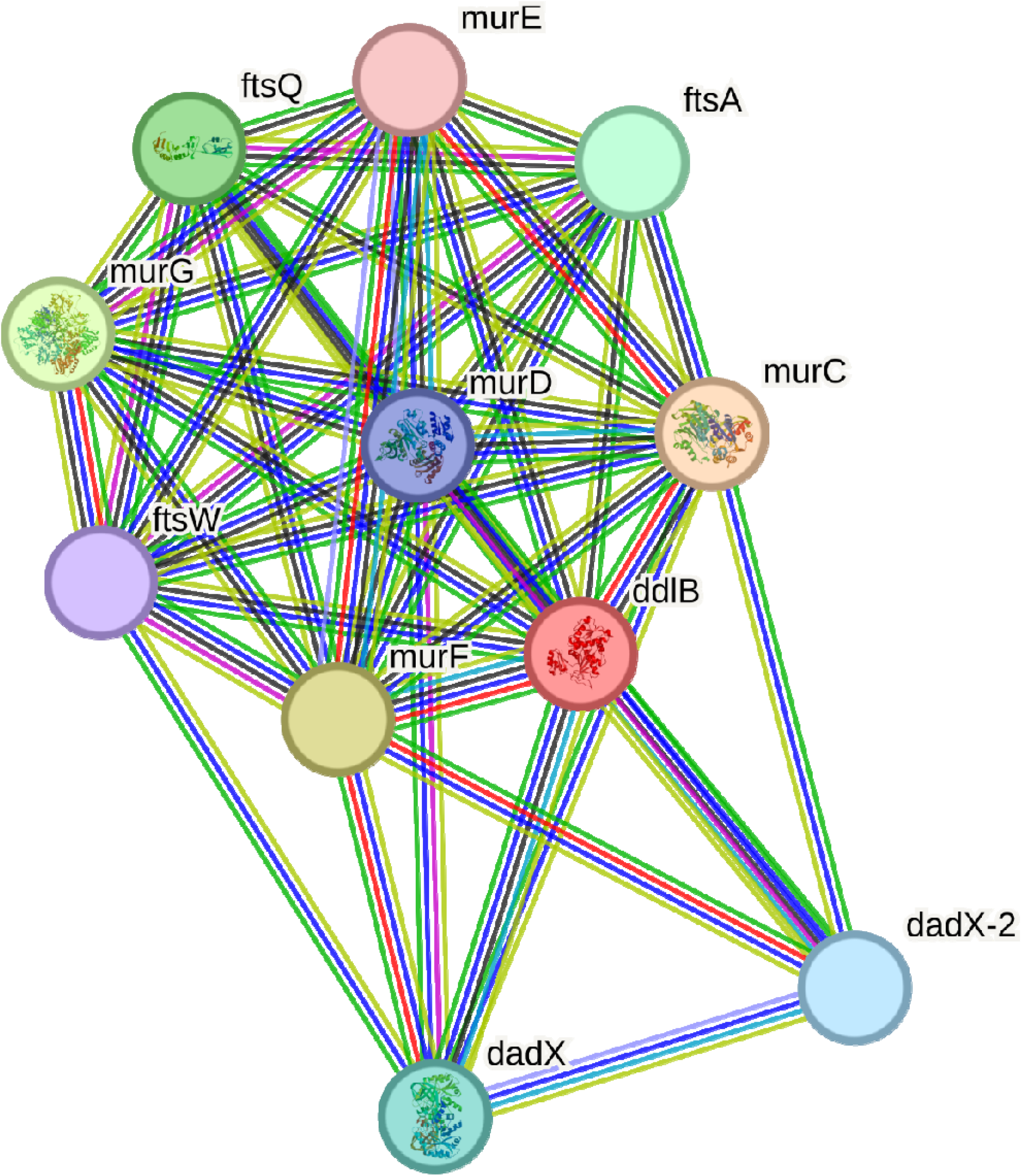
The result from String server where the bacterial protein D-Alanine-D-Alanine Ligase interacts with a variety of proteins that are responsible for cell wall biosynthesis

### Brute force screening

Brute force screening against BpDdl was performed by targeting the D-alanine-D-alanine binding pocket using molecular docking. Total 5035 compounds were screened, using AutoDock Vina (Morris et al., 2009a) through PyRx (Dallakyan & Olson, 2015) platform. Among them only top 4 compounds were selected and the binding pose of those compounds in the binding pocket were validated using the Chimera software (Pettersen et al., 2004). The AutoDock Vina score (ADV) scores in terms of binding energy of the top four compounds from higher to lower are tabulated in Table 1 with their chemical structures. From the results it was observed that the average binding affinity across all four drugs is -10.05 kcal/mol. Rimegepant (RIM) exhibited the highest binding affinity with an ADV score of –10.8 kcal/mol and Radotinib (RAD) exhibited the second highest binding affinity of –10.7 kcal/mol. Both Conivaptan (CON) and Nilotinib (NIL) exhibited near average binding affinity i.e. -10.3 kcal/mol. The amino acids residues present in the binding cavity of BpDdl interacting with the top four ligands (RIM, RAD, CON, NIL) are represented in Figure 5. The contribution of hydrogen bond and hydrophobic interaction in the binding of the top four ligands were determined using LIGPLOT+ (Figure 6) (Wallace et al., 1995),where the hydrogen bonds are represented as green dotted line and hydrophobic interactions are represented as red spoked arcs. RIM exhibited the highest binding energy is attributed to the six-hydrogen bonding interaction that takes place with the Tyr221, Arg260, Asn277, Lys220 and Ser286 residues (Figure 6A). The Ser286 residues is used to form two hydrogen bonds with the drug. RAD is attributed to the three-hydrogen bonding that takes place between the drug and the His285, Ala223 and Ser286 residues of target protein (Figure 6B). CON forms two hydrogen bonds with Lys104 and Asn277 residues of the target protein (Figure 6C) whereas NIL forms two hydrogen bonds with Tyr215 and Asn277 residues (Figure 6D). These top four compounds that showed significantly stronger binding affinity were subjected to ADMET analysis and toxicity prediction (Table 2) followed by Molecular Dynamics (MD) simulation.

**Figure 5:**
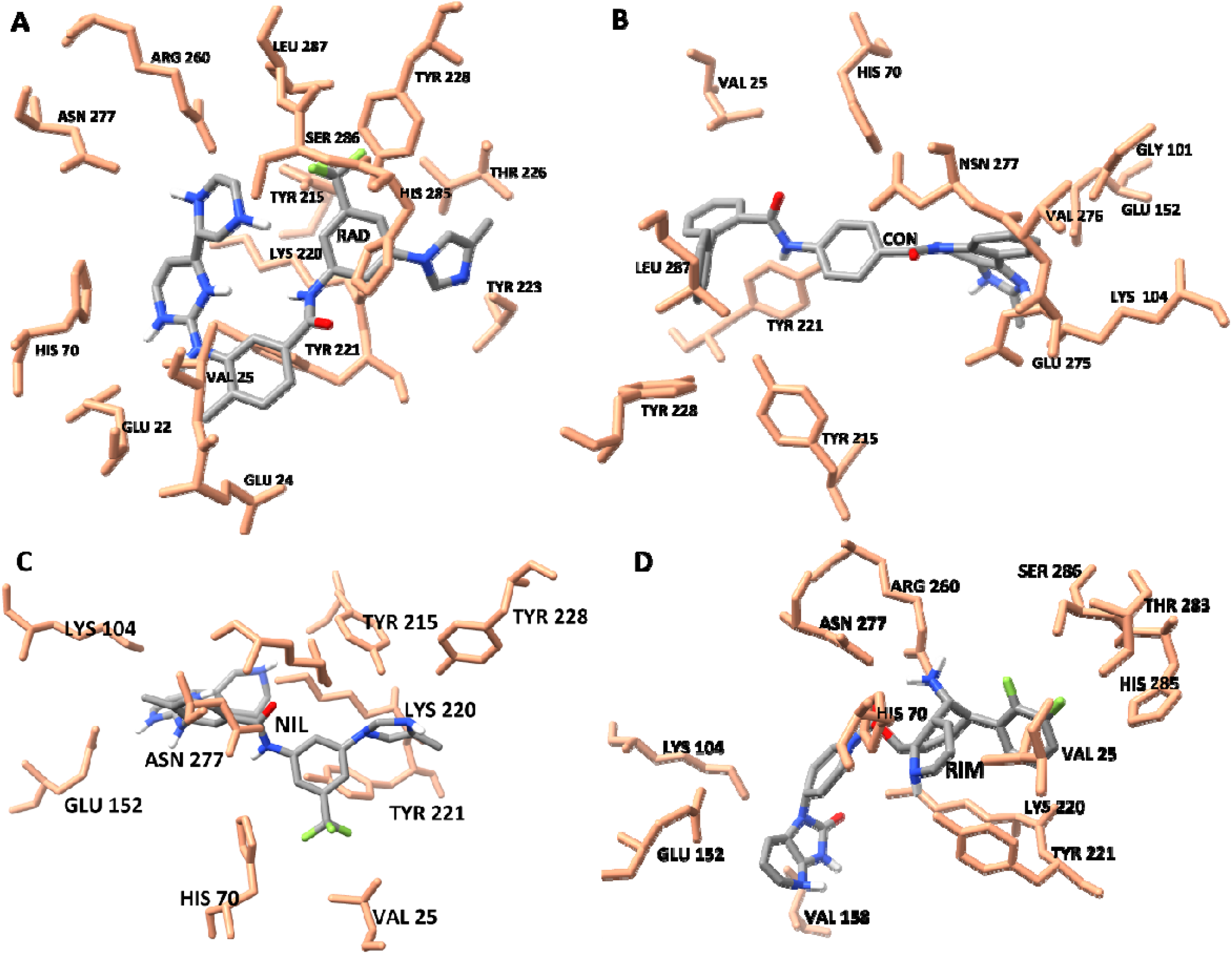
The binding poses and interacting amino acid residues of (A) RAD (B) CON (C) NIL (D) RIM with BpDdl and the interacting residues are shown in sticks and labelled with the three-letter code.

**Figure 6:**
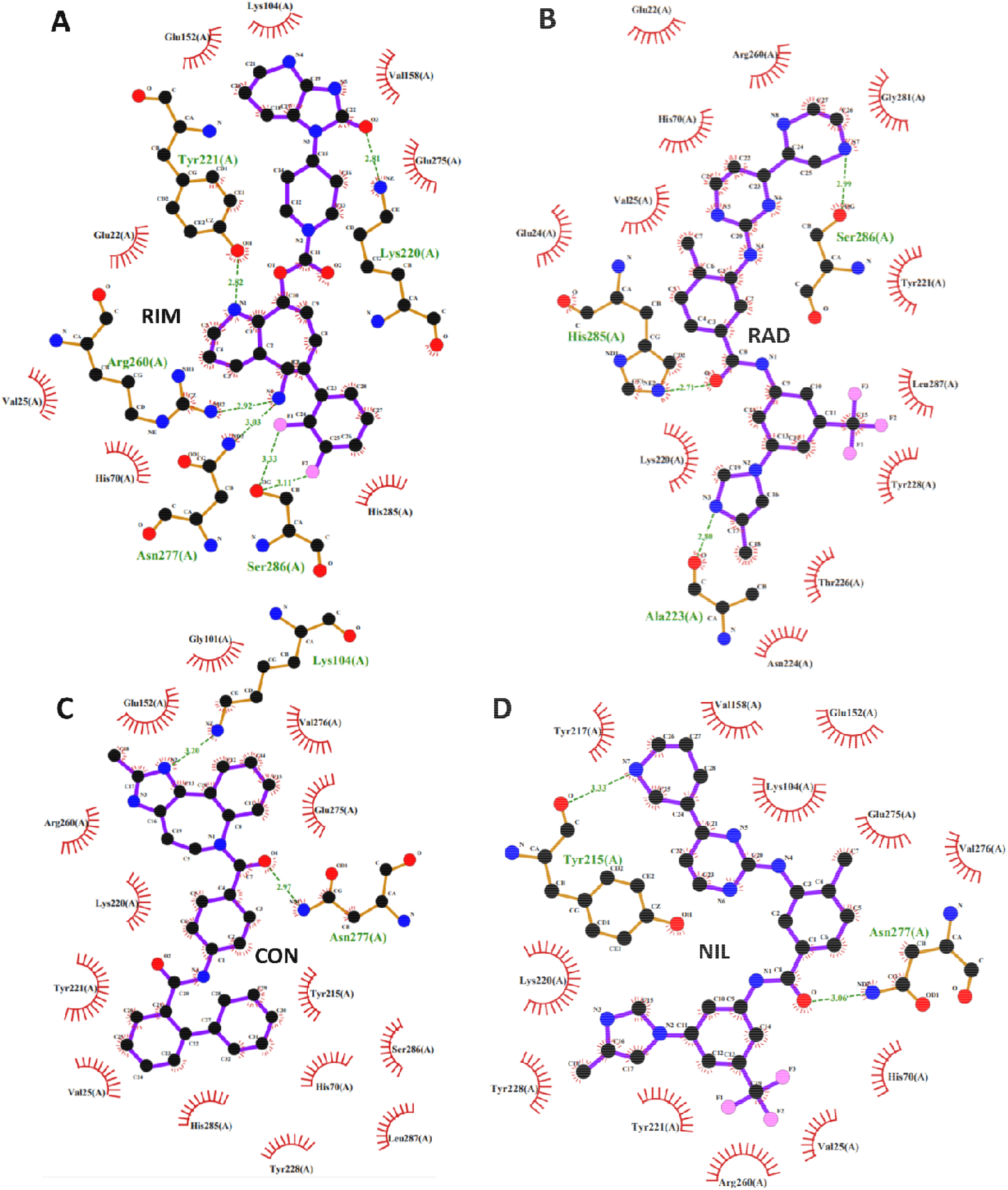
Visual representation of hydrogen bond and hydrophobic interactions of (A) RIM (B) RAD (C) CON (D) NIL with BpDdl, where the green dotted line indicates the hydrogen bond and spoked arcs indicates the hydrophobic interactions.

**Table 1:**
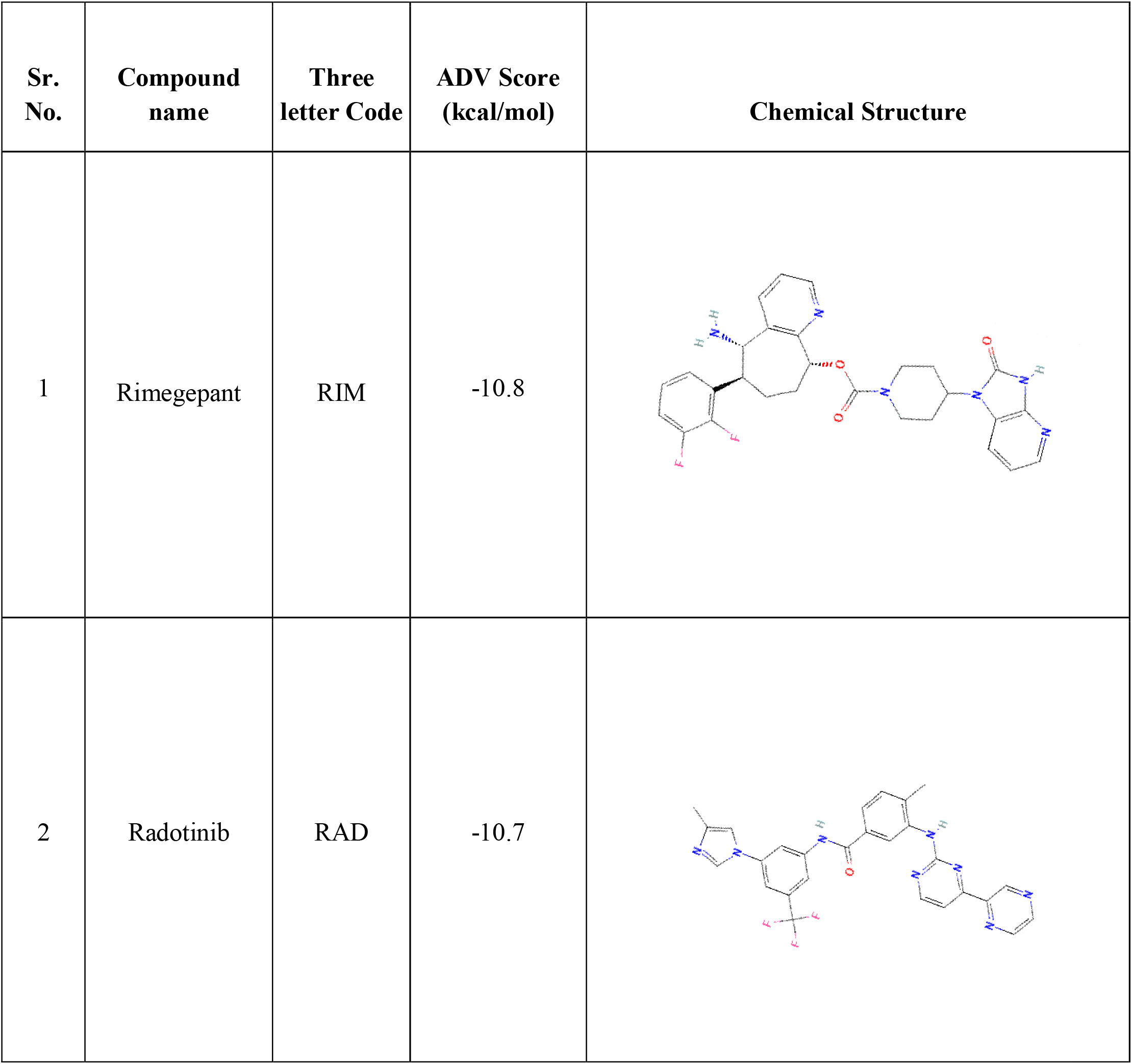
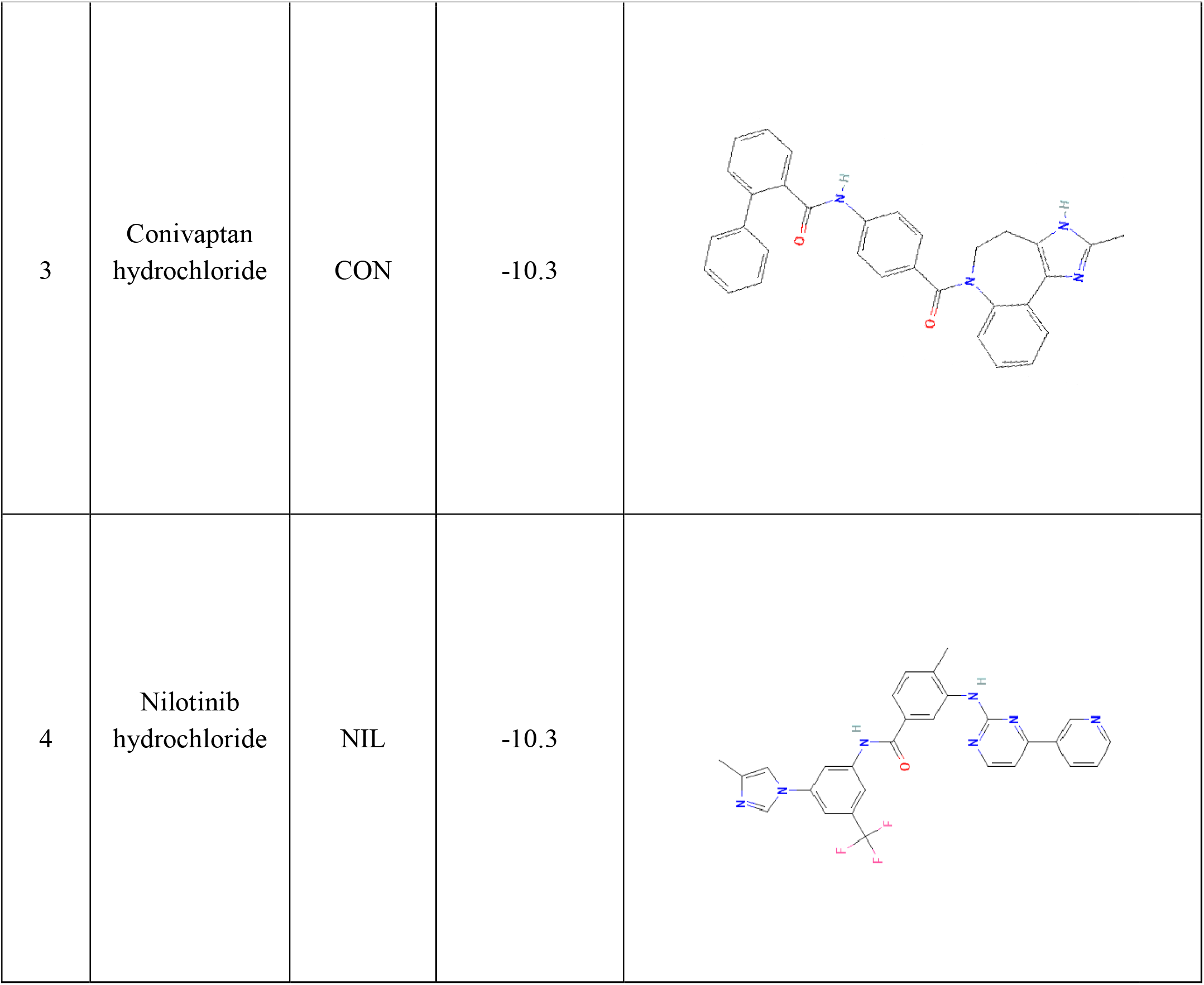

**Table 2:** ADME and toxicity analysis.

| Parameters | RIM | CON | RAD | NIL |
| --- | --- | --- | --- | --- |
| **MW** | 534.56 | 535.04 | 530.5 | 565.98 |
| **Acceptable range** | ≤ 500 | ≤ 500 | ≤ 500 | ≤ 500 |
| **NHBA** | 8 | 3 | 9 | 8 |
| **Acceptable range** | ≤ 10 | ≤ 10 | ≤ 10 | ≤ 10 |
| **NHBD** | 2 | 2 | 2 | 2 |
| **Acceptable range** | ≤ 5 | ≤ 5 | ≤ 5 | ≤ 5 |
| **MR** | 143.49 | 160.51 | 138.87 | 148.04 |
| **Acceptable range** | 40–130 | 40–130 | 40–130 | 40–130 |
| **Mlogp** | 2.59 | 4.26 | 1.78 | 2.94 |
| **Acceptable range** | ≤ 4.15 | ≤ 4.15 | ≤ 4.15 | ≤ 4.15 |
| **ProTox class** | 4 | 4 | 4 | 4 |

### Molecular Dynamics Simulation

The top four compounds based on ADV energy that passed Lipinski’s Rules, underwent molecular dynamics (MD) simulation, followed by binding free energy calculations to conduct a second round of screening. This process aimed to identify stronger interactions among the top four compounds. Key metrics including root mean square deviation (RMSD) (Schreiner et al., 2012), root mean square fluctuation (RMSF), and radius of gyration (R_g_) (Lobanov et al., 2008) were calculated from the MD simulation trajectories. The average RMSD, RMSD, R_g_ and ligand RMSD has been tabulated in Table 3.

**Table 3:** Molecular Dynamics simulation matrics.

| SL NO. | Compound code | Average RMSD (nm) | Average Ligand RMSD (nm) | Average R<sub>g</sub> (nm) | Average RMSF (nm) |
| --- | --- | --- | --- | --- | --- |
| 1 | RIM | 0.236 | 0.174 | 2.039 | 0.087 |
| 2 | RAD | 0.184 | 0.574 | 2.044 | 0.101 |
| 3 | CON | 0.238 | 0.478 | 2.088 | 0.119 |
| 4 | NIL | 0.188 | 0.292 | 2.030 | 0.089 |

The average RMSD for the CON_Ddl complex is 0.237 nm, NIL_Ddl complex is 0.1875 nm and RIM_Ddl is 0.2360 nm (Table 3). The RMSD values of drug bound protein complex is much lower compared to the apo protein with an RMSD value of 0.327 (Figure 7A). The RMSF of the protein alone is 0.1472 nm, while the RMSF for protein-inhibitor complexes ranges from 0.0867 nm to 0.1190 nm (Figure 7B). The radius of gyration (R_g_) indicates the compactness of the protein. The average R_g_ value for the protein is 2.155 nm. The R_g_ values for protein-inhibitor complexes range from 2.030 nm to 2.088 nm (Figure 7C). The ligand RMSD of the top four protein complexes were calculated from the MD simulation and represented in Figure 7D.

**Figure 7:**
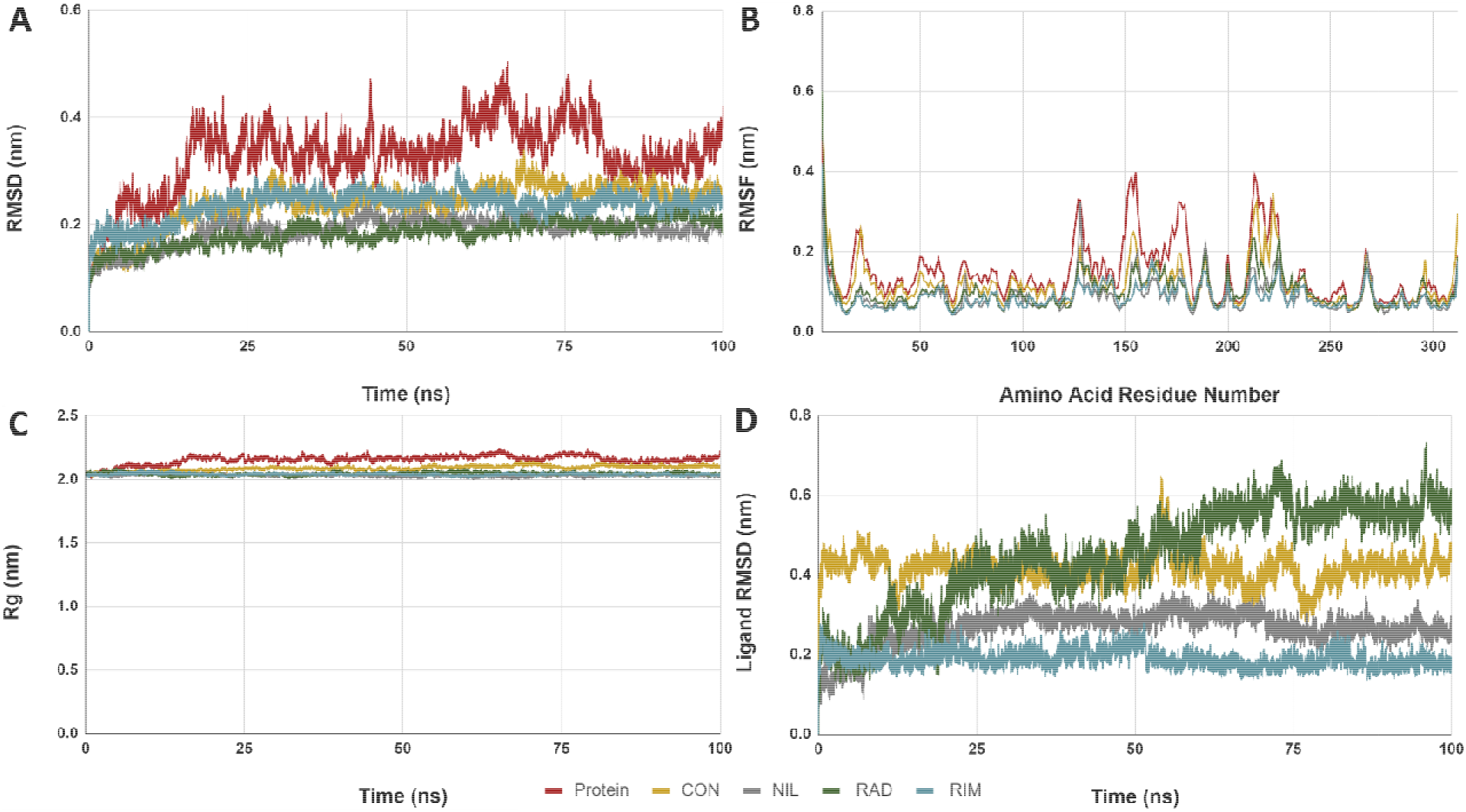
Molecular dynamics metrics of BpDdl with top inhibitors. The inhibitor bound BpDdl complexes and the protein alone are shown in different colours. (A) The RMSD (nm) of backbone vs time in ns (B) backbone RMSF (nm) against the amino residue number, (C) Radius of gyration (R_g_) in nm vs time (ns) and (D) Ligand RMSD (nm) of top four inhibitors vs time (ns)

### Principal Component Analysis

Principal Component Analysis (PCA) was done to observe the collective motion of Ddl over a 100 ns simulation, focusing on Cα atoms. The Eigenvalue Rank plots demonstrated that PC1 and PC2 encapsulated most of the collective motion. Scatter dot plots for PC1 and PC2 values for each protein across all dynamic systems, with each dot representing the protein’s collective motion (Cα atoms) at different simulation time points (blue: initial, white: intermediate, red: final). Dots clustering around the center (0, 0) indicated less motion and a stable conformation, while scattered dots indicated greater motion and structural instability. For BpDdl-conivaptan, BpDdl-nilotinib and BpDdl-radotinib complex, the blue/white/red dots are clustering in a narrower space compared to the apo form of the protein that indicated that the target protein is stabilized with the binding of the drug (Figure 8). Protein forming complex with Rimegepant is forming cluster around the center but with a wider space than the apo protein.

**Figure 8:**
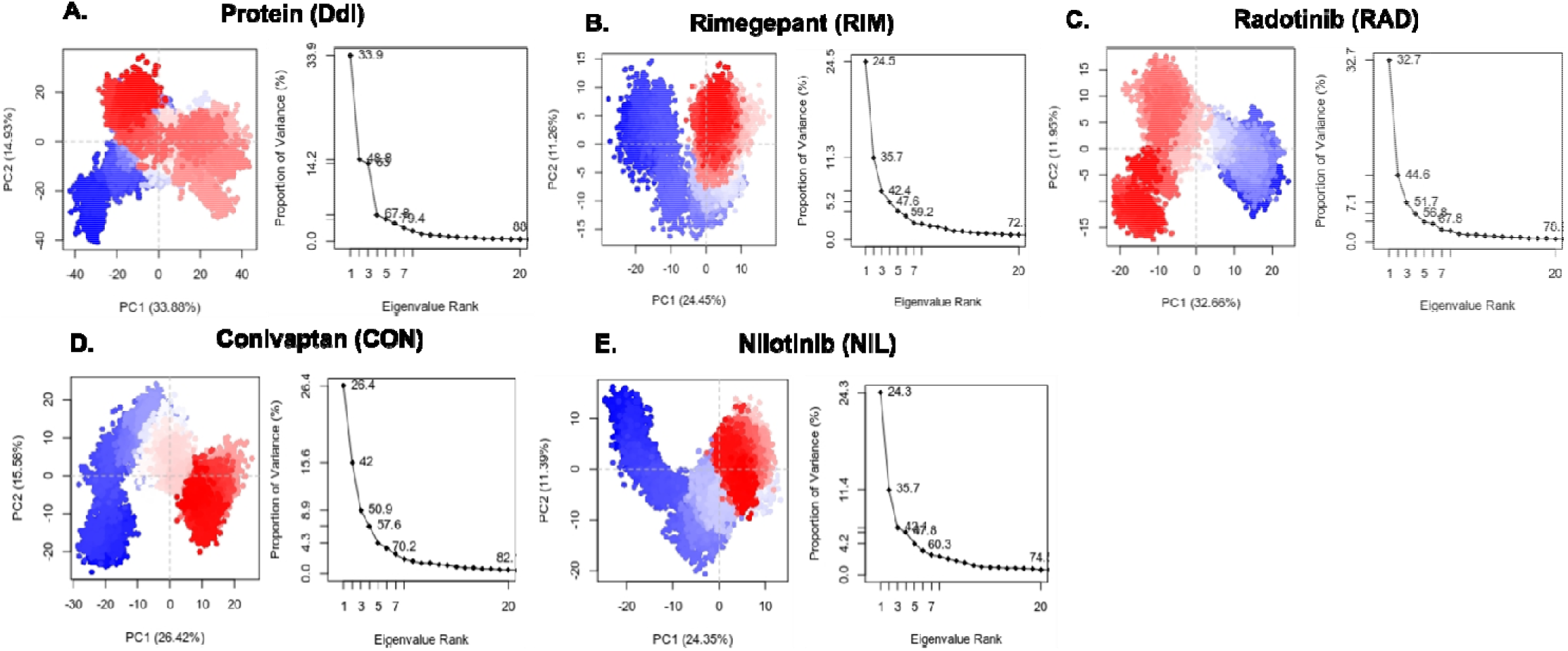
PCA of the BpDdl alone and complexed with top four inhibitors by using eigenvector 1 eigenvector 2. (A) BpDdl (B) RIM (C) RAD (D) CON (E) NIL, blue dots represent the initial timescale and red dots represent final timescale.

### Dynamic Cross-Correlation Matrix (DCCM) Analysis

Dynamic cross-correlation matrix was created using the coordinates of Cα atoms from each dynamic system to analyze the comparative motion of amino acid residues. The residue motions can be positively (cyan), negatively (pink), or neutrally (white) correlated. The findings revealed that the residues in the Apo-form BpDdl, as well as in ligand bound form exhibited highly correlated motions (Figure 9).

**Figure 9:**
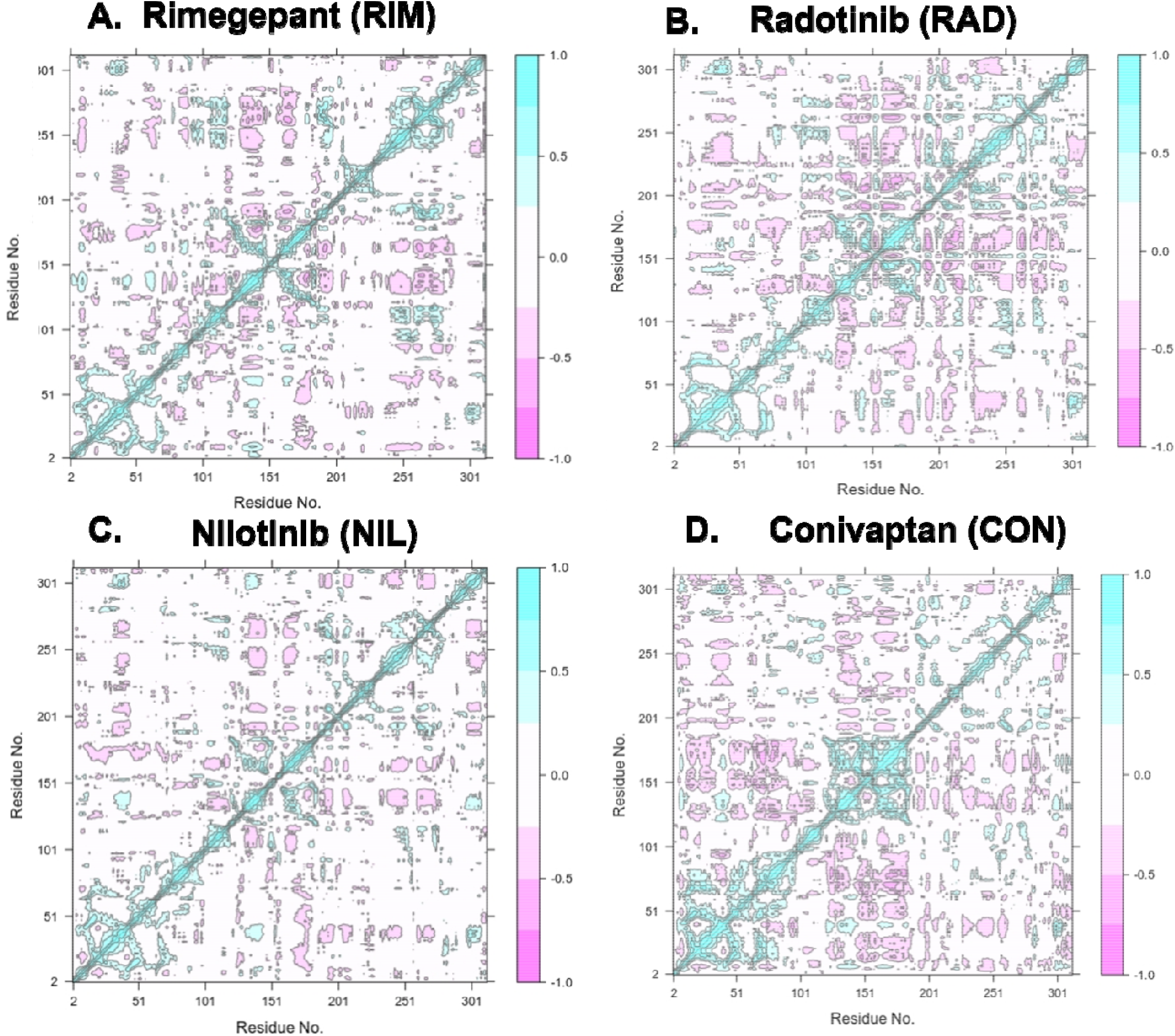
Dynamic cross corelation matrix of Cα atoms of BpDdl-inhibitor complexes (A) RIM (B) RAD (C) NIL (D) CON. In the plot the positively corelated residues are shown in cyan, negatively correlated residues are in pink and neutral are shown in white colour.

### Binding Free Energy

The difference in the binding energy depends on various factors like Van Der Waals Forces, Electrostatic Interactions and Non-Bonding Forces. The highest binding energy was exhibited by the BpDdl-Conivaptan complex with a binding energy of -66.45 +/-3.23 kcal/mol and the BpDdl-Rimegepant complex with a second highest binding energy of -60.63 +/-2.54 kcal/mol. The lowest binding energy was exhibited by the BpDdl-Radotinib complex with a binding energy of -8.0+/-3.4 kcal/mol. An intermediate binding affinity was exhibited by BpDdl-Nilotinib complex with a binding affinity of -37.17 +/-3.22 kcal/mol. The binding affinity of different protein-ligand complexes are described in the Figure 10.

**Figure 10:**
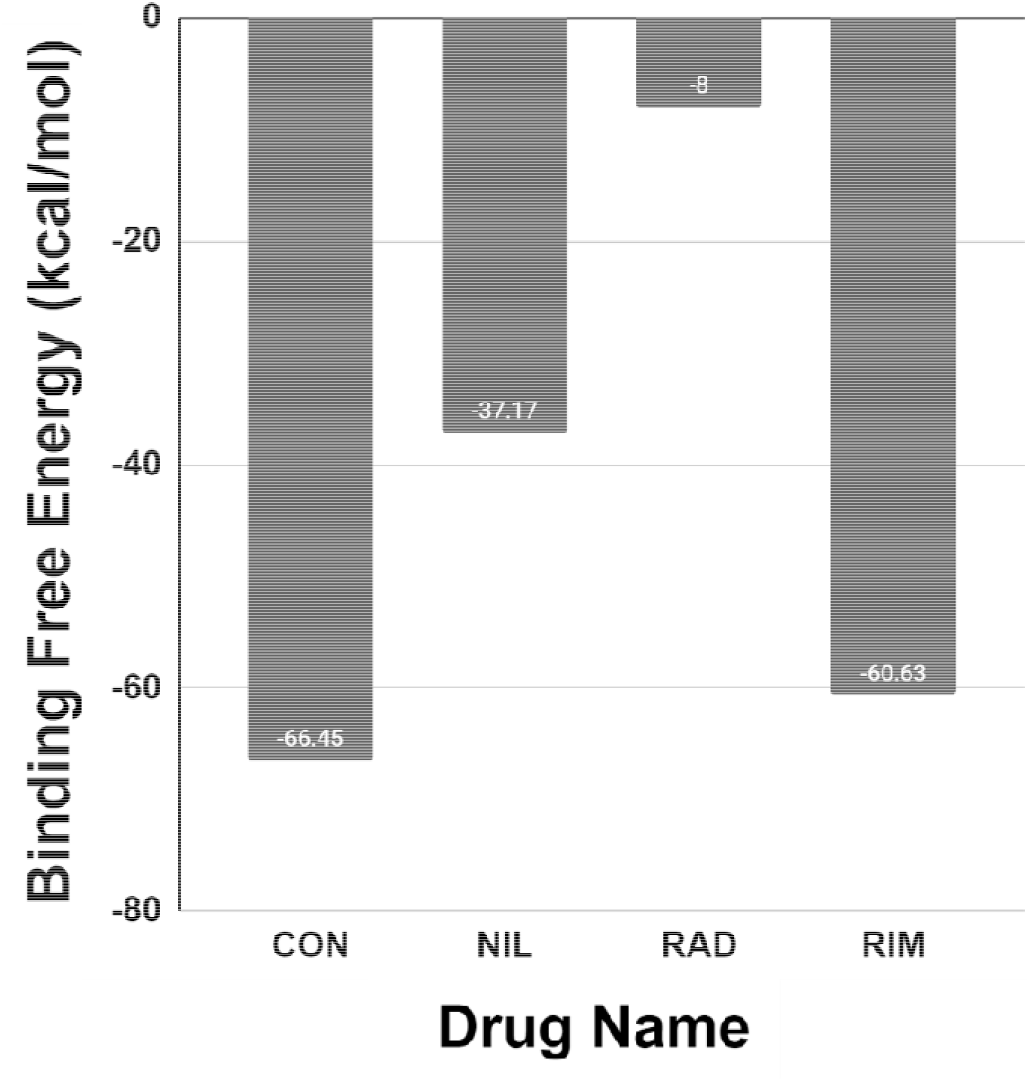
Graphical representation of binding free energy (ΔG) for top four, BpDdl-inhibitor complex expressed in kJ/mol.

### Free Energy Landscape

The free energy landscape (FEL) is a valuable tool for examining complex biological systems. It maps the energy of a system onto a multidimensional landscape, where each dimension represents a specific structural degree of freedom. This approach offers a detailed view of the system’s energetics, considering both internal energy and entropy. This process uses the first two principal components (PC1 and PC2) as input to plot the Gibb’s free energy landscape by inverting a multidimensional histogram using the Boltzmann formula.

Visualizing the FEL provides insights into the structural and energetic characteristics of the system. The global minimum on the FEL indicates the most energetically favorable structural configuration, while local minima represent other energetically stable conformations. Understanding the FEL is crucial for studying protein stability, folding, function, and predicting the behavior of drug molecules. Figure 11 describes Gibb’s free energy landscape of the top three compounds and the apo protein.

**Figure 11:**
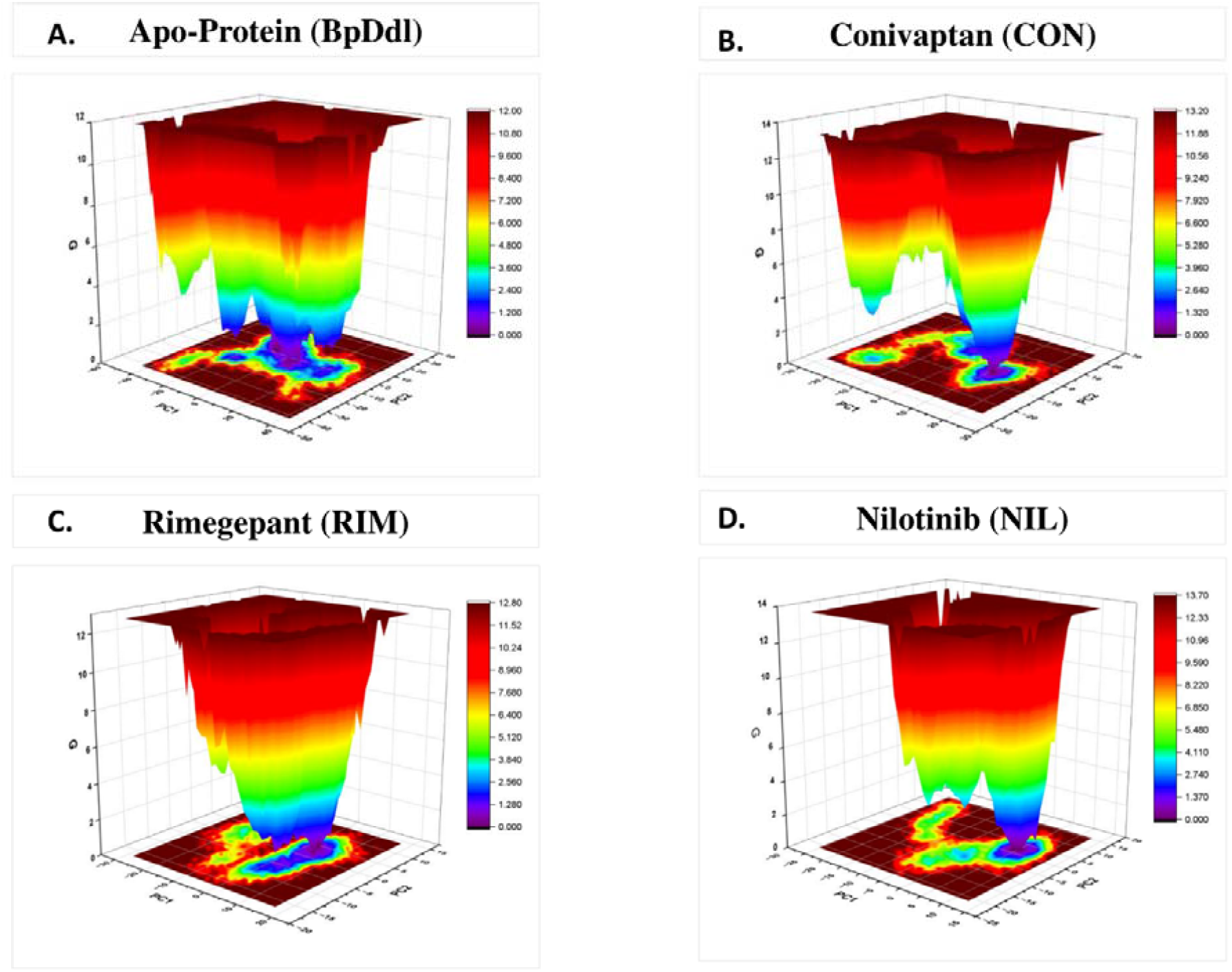
The free energy landscape (FEL) of BpDdl with its inhibitor bound form represented as the colour coded scale. (A) Apo-protein BpDdl (B) CON (C) RIM (D) NIL. The ranges of colour scale in FEL graph has shown from dark blue/violet to red/yellow indicates lower energy to higher energy, whereas green indicates the intermediate energy states.

## DISCUSSION

In this study, the bacterial protein D-Alanine-D-Alanine Ligase (Ddl) was used as a potential drug target against *Burkholderia pseudomallei* strain K96243. D-Alanine-D-Alanine Ligase (Ddl) is an enzyme essential for peptidoglycan synthesis of bacterial cell wall and this enzyme has acted as a drug target in other organisms as well (Bruning et al., 2011).

The Ddl in *Burkholderia pseudomallei* strain is a 312 amino acid protein with molecular weight of 33.34 kD. It is a member of the ATP-grasp superfamily, exhibiting a highly conserved fold consisting of three well-defined domains: N-terminal (65aa-95aa), central, and C-terminal (110aa-307aa). Specifically, in Ddl, the secondary structure includes 10 α-helices, and 12 β-strands.

The N-terminal segment forms an α/β-domain, while the central and C-terminal domains feature the ATP-grasp fold, comprising two α/β-domains that create a narrow pocket for ATP binding. The D-Ala binding part of the active site is in the C-terminal domain, with the P-loop (residues 149–159) in the central domain interacting with the γ-phosphates of ATP, and the π-helix configuration of the x-loop (residues 198–222) in the C-terminal domain. Notable structural differences are observed between subunits of the two BpDdl structures, particularly in the central domains and several helices in the N-terminal and C-terminal domains, which adopt different positions lining the active site.

The presence of isoform B of Ddl in *Burkholderia pseudomallei* instead of the cooperative activity of both Isoform A and B in the case of pathogenic *Enterobactericeace* and *Salmonella typhimurium* confers a significant weak point in the bacterial defenses that is attempted to exploit in our study. After the target protein was selected, it was subjected to extensive structural and evolutionary analysis. Protein Blast results indicate that BpDdl shares significant sequence identity and similarity with the Ddl protein of other pathogens like *Mycobacterium tuberculosis, Staphylococcus aureus, Streptococcus pneumoniae*, etc. thus the drug can be used against other pathogens as well. The drugs that were identified from molecular docking were subjected to Molecular Dynamic Simulations (MDS) where the root mean square deviation (RMSD), root mean square fluctuations (RMSF) and radius of gyration (R_g_) were calculated. The RMSD values exhibited that the drugs are forming stable complexes with the protein since the RMSD of the complex is lower than the RMSD value of apo-protein. The stability of the drug ligand complex was also confirmed by the RMSF and R_g_ values. The binding affinity analysis by the MMPBSA revealed that conivaptan exhibited the highest binding affinity with a free energy of -66.45 +/-3.23 kcal/mol and rimegepant exhibited the second highest free energy of -60.63 +/-2.54 kcal/mol. Analysing the data from PCA and FEL it is evident that conivaptan is a suitable inhibitor against BpDdl. Rimegepant also has binding affinity close to conivaptan but the lower complex stability makes it less ideal for drug inhibition. Conivaptan is a vasopressin receptor antagonist used primarily for the treatment of euvolemic and hypervolemic hyponatremia. It acts by blocking both V1a and V2 receptors of vasopressin, thereby promoting free water excretion without significant loss of electrolytes (THIBONNIER, 2003). This pharmacological action helps in correcting hyponatremia by increasing serum sodium levels in patients with various underlying conditions such as congestive heart failure, liver cirrhosis, and the syndrome of inappropriate antidiuretic hormone secretion (SIADH) (Cada et al., 2006).

## CONCLUSION

In conclusion, our study provides evidence supporting the potential use of Conivaptan as a specific D-Alanine-D-Alanine Ligase inhibitor to disrupt cell wall formation and expose the pathogen to the hostile host cellular environment. The findings underscore the importance of specificity in BpDdl inhibition and offer valuable insights into developing novel therapeutic strategies against Mellidosis (Whitmore Disease). However, a comprehensive understanding of Conivaptan’s impact on the pathogen and its implications for anti-bacterial activity requires further in-vitro studies. The current research does not fully explore these complex dynamics.

## MATERIALS AND METHODS

### Identification of Metabolic Pathways

The metabolic pathways of *Burkholderia pseudomallei* K96243 (KEGG organism: bps) and the human host (KEGG organism: hsa) were obtained from the Kyoto Encyclopedia of Genes and Genomes (KEGG; https://www.genome.jp/kegg/) (Kanehisa et al., 2021). Subsequently, these pathways were manually compared to identify unique and common features. The proteins responsible for driving these pathways were then extracted from the Uniprot database (https://www.uniprot.org) (“UniProt: A Worldwide Hub of Protein Knowledge,” 2019) and the NCBI database (RefSeq ID: https://www.ncbi.nlm.nih.gov/) (O’Leary et al., 2016). This comparative analysis allowed for the identification of metabolic pathways and associated proteins in both the pathogen and the human host, aiding in the understanding of potential targets for therapeutic intervention.

### Assessment of Essential Genes

The Database of Essential Gene (DEG; http://origin.tubic.org/deg/public/index.php/index) (Luo et al., 2014) holds the information of the essential genes i.e. the genes that drive life in archea, bacteria and eukarya. Presently this database houses information on bacterial genes (Updated on). The essential genes of the K96243 were identified by integrating the results of BlastP analysis and DEG database. Only the proteins that passed the criteria of e-value of <0.0001 and the bit score of >100 were used for further analysis (Hajduk & Greer, 2007).

### Identification of Non-Homologous Genes

The identification of non-homologous genes, which are present in the pathogen but absent in the host, is crucial for drug design to avoid nonspecific targeting of human proteins, which may lead to side effects and cross-reactivity (Omelchenko et al., 2010). To achieve this, protein sequences were subjected to the NCBI BlastP server (https://blast.ncbi.nlm.nih.gov/Blast.cgi). Subsequently, a similarity search was conducted between the proteins of Burkholderia pseudomallei K96243 and human proteins. Only proteins with an e-value <0.005 and a sequence similarity <35% were selected for non-homology analysis (Pearson, 1995). This approach ensured the identification of pathogen-specific targets for drug development while minimizing potential off-target effects in the host.

### Identification of Subcellular Localization of Target Proteins

Following established research norms, proteins located in the extracellular space, periplasm, and cell surface are typically targeted for vaccine development, while proteins found in the inner membrane and cytosol are selected as drug targets. All target proteins obtained were assessed using PSORTb 3.0 (https://www.psort.org/psortb/) (N. Y. Yu et al., 2010) and CELLO v2.5 (http://cello.life.nctu.edu.tw/) (C. Yu et al., 2004) to determine their subcellular localization. Proteins annotated as extracellular were further analyzed for the presence of transmembrane domains using the TMHMM v2.0 server (http://www.cbs.dtu.dk/services/TMHMM/) (Krogh et al., 2001). Proteins identified as extracellular, or cell surface were then chosen for additional antigenicity testing. This systematic approach ensured the selection of suitable vaccine targets based on their cellular localization and potential immunogenicity.

### Analysis of Immunogenicity of Cell Surface Proteins

In vaccine design, evaluating the immunogenic potential of target proteins is paramount. To accomplish this, the VaxiJen v2.0 server (http://www.ddg-pharmfac.net/vaxijen/VaxiJen.html) (Doytchinova & Flower, 2007) was utilized. Proteins demonstrating an antigenic potential above 4.0 were selected for further analysis. These proteins, meeting the threshold criteria, were then chosen for epitope mapping, enabling the identification of specific regions that could elicit an immune response. This process ensured the selection of promising vaccine candidates with high antigenic potential.

### Potential of the Non Homologous Proteins as Drug Targets

In accordance with prior research, the druggability of proteins can be assessed by evaluating the binding affinity of similar proteins to drug-like molecules (Hajduk et al., 2005). Potential drug targets were identified by querying against the DrugBank Database (https://go.drugbank.com/) (Wishart et al., 2018) and the Selleckchem Database (https://www.selleckchem.com/), which contains FDA-approved drugs, utilizing the inbuilt BLAST tool. All non-homologous protein targets were analyzed with default search parameters (i.e., e-value <0.0001). This approach facilitated the identification of proteins with potential drug-binding capabilities, aiding in the selection of viable drug targets for further investigation.

### Protein-Protein Interaction

Understanding protein-protein interactions is crucial in drug design and development, as they determine the efficacy and potential of a drug. Essential proteins, which are vital for the survival of an organism, typically have numerous interacting partners that contribute to downstream signaling mechanisms (Nikolaidis et al., 2014). By disrupting these pathways with inhibitors, it is possible to interfere with the normal metabolic processes of the pathogen, ultimately leading to its eradication. To investigate these pathways involving essential proteins, the STRING database (https://string-db.org/) (Szklarczyk et al., 2017) was utilized. This platform enabled the mapping of interactions between non-homologous target proteins, thereby creating an intra-species protein-protein interaction network. Through this analysis, insights into the complex molecular networks within the pathogen were gained, facilitating the identification of potential drug targets and the development of targeted therapeutic strategies.

### Crystal Structure Selection and Fixing Missing Residues

For the study, the protein D-alanine-D-alanine ligase (UniProt ID: A3NZL3; PDB ID: 5NRI) was selected. This cytoplasmic protein primarily facilitates cell wall formation and maintenance through peptidoglycan biogenesis. Its significance extends beyond bacterial protection and sustenance to include a pivotal role in antibiotic resistance (Larsson & Flach, 2022). The suitability of targeting this protein lies in the fact that bacterial cells cannot survive within hosts without a functional cell wall, rendering them incapable of developing antibiotic resistance mechanisms. The crystal structure of D-alanine-D-alanine ligase (PDB: 5NRI) was obtained from the RCSB PDB (https://www.rcsb.org/structure/5NRI). Missing structural information was addressed using computational modeling tools such as Robetta, Phyre2, and SWISS-MODEL Workspace (Waterhouse et al., 2018). The quality of the resulting model was assessed and validated using QMEAN (Qualitative Model Energy Analysis) (https://swissmodel.expasy.org/), ProSA (https://prosa.services.came.sbg.ac.at/prosa.php) (Wiederstein & Sippl, 2007), and the PROCHECK server (https://www.ebi.ac.uk/thornton-srv/software/PROCHECK/) (Laskowski et al., 1993). Furthermore, the model underwent optimization through a 200-step steepest descent energy minimization process using the Swiss PDB Viewer (https://spdbv.unil.ch/).

These steps ensured the refinement and accuracy of the structural model, providing a reliable basis for subsequent analyses and drug design endeavors.

### Structure-Based Virtual Screening

The active site of D-alanine-D-alanine ligase was scrutinized and targeted for drug identification. Small molecule library was prepared from SelleckChem and DrugBank. The receptor structure was prepared for virtual screening by adding polar hydrogen atoms and Kollman charges. A total of 3105 FDA-approved drugs were obtained from the DrugBank Database and 1930 FDA-approved drugs were obtained from Sellekchem Database in SDF file format for screening. Ligand preparation utilized the prepareligand4.py python script provided by AutoDock developers (Morris et al., 2009a), with Open Babel used to convert all drug compound files into PDBQT format (O’Boyle et al., 2011). A receptor grid-box was generated with dimensions of 17 Å × 17 Å × 20 Å, centered at x = 46, y = 25, z = 74. Virtual screening of processed drugs against D-alanine-D-alanine ligase was performed using AutoDock Vina (Trott & Olson, 2010). Subsequently, the top ten drugs obtained from virtual screening were subjected to docking analysis using AutoDock 4.2 (Morris et al., 2009a, 2009b).

### Molecular Docking Analysis

The receptor, D-alanine-D-alanine ligase underwent pre-processing for docking in AutoDock 4.2 (Morris et al., 2009a). Polar hydrogens were added to the receptor followed by the assignment of Kollman charges. Non-polar hydrogens were merged, and Gasteiger charges were assigned to the selected ligands using AutoDock 4.2. Molecular docking was conducted using the same grid box parameters: dimensions of 17 Å × 17 Å × 20 Å, centered at x = 48, y = 26, z = 75. Lamarckian Genetic Algorithm (GA) was employed for a total of 10 runs. AutoDock calculated the free energy of binding (ΔG), considering intermolecular energy components (vdW, H-bond, desolv Energy, Electrostatic Energy), internal energy, and torsional energy, subtracting the unbound system’s internal energy (Morris et al., 2009a). The inhibition constant (Ki) value was derived using the equation [Ki = exp(ΔG/(R^*^T)]. The docking results were visualized using ChimeraX v1.15 (Pettersen et al., 2004), and receptor-ligand interactions were manually analyzed with LigPlot+ (Laskowski & Swindells, 2011) to elucidate their molecular interactions.

### Drug-Like Properties And Prediction Of Toxicity / Pharmacokinetics

The top 10 compounds that were chosen based on the ADV energy were subjected to SwissADME server for screening of ADMET (Absorbtion, Digestion, Metabolism, Excretion) properties and ProTox II server was used to check the toxicity scores.

### Molecular Dynamic Simulation

The preliminary selection of drug molecules based on their AutoDock Vina (ADV) energy within the protein-ligand complex was followed by Molecular Dynamics Simulation (MD Simulation) and subsequent Molecular Mechanics Poisson-Boltzmann Surface Area (MMG BSA) calculations. All MD simulations were executed on the GROMACS 5.1.2 platform (http://www.gromacs.org/) employing the CHARMM 36 force field (Vanommeslaeghe et al., 2010). The protein-ligand complex was solvated using the TIP3P water model (Jorgensen et al., 1983) within a cubic box. The topology file of the solvated and electroneutral complex underwent energy minimization utilizing the steepest descent algorithm while maintaining forces below 10.0 kJ/mol. Prior to the final MD run, the system underwent equilibration through two stages: NVT (isothermal-isochoric ensemble) and NPT (isothermal-isobaric ensemble). In the NVT stage, the temperature was maintained at 300 K for 100 ps using the Berendsen method (Berendsen et al., 1984). Subsequently, in the NPT stage, pressure was held constant at 1 bar while the number of particles, pressure, and temperature were equilibrated for 100 ps using the Rahman barostat method (Parrinello & Rahman, 1981). Following equilibration, the system underwent MD simulation. Trajectories from MD simulations were corrected for periodic boundary conditions and centered. From the corrected trajectories, Root Mean Square Deviation (RMSD) (Schreiner et al., 2012), Root Mean Square Fluctuation (RMSF) (Maiorov & Crippen, 1994), and Radius of Gyration (R_g_) (Lobanov et al., 2008), were calculated. The last 10 ns of the corrected trajectory from each MD simulation experiment were extracted to compute the protein-ligand binding free energy (ΔG_binding) using the MM-PBSA method with the g_mmpbsa script (Kumari et al., 2014). The binding free energy was calculated using the following equation where G_Complex_ indicates the binding free energy of the protein-inhibitor complex, G_Receptor_ refers to the free energy of receptor and G_Ligand_ represents the free energy of ligand:

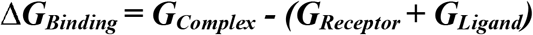

