## Supplementary table 1 for "Subtractive proteomics unravel the potency of D-Alanine-D-Alanine Ligase as the drug target for *Burkholderia pseudomallei*"

**Supplementary table 1:** Unique pathways (KEGG) for *Burkholderia pseudomallei* K96243 (TaxID: 272560) compared to Homo sapiens (TaxID: 9606)

| **Sl No.** | **Unique pathways (KEGG) for B. pseudomallei K96243** | **KEGG pathway ID** |
| --- | --- | --- |
| 1 | Monobactam biosynthesis | [bps00261](https://www.kegg.jp/entry/bps00261) |
| 2 | Geraniol degradation | [bps00281](https://www.kegg.jp/entry/bps00281) |
| 3 | Lysine biosynthesis | [bps00300](https://www.kegg.jp/entry/bps00300) |
| 4 | Penicillin and cephalosporin biosynthesis | [bps00311](https://www.kegg.jp/entry/bps00311) |
| 5 | Carbapenem biosynthesis | [bps00332](https://www.kegg.jp/entry/bps00332) |
| 6 | Chlorocyclohexane and chlorobenzene degradation | [bps00361](https://www.kegg.jp/entry/bps00361) |
| 7 | Benzoate degradation | [bps00362](https://www.kegg.jp/entry/bps00362) |
| 8 | Fluorobenzoate degradation | [bps00364](https://www.kegg.jp/entry/bps00364) |
| 9 | Novobiocin biosynthesis | [bps00401](https://www.kegg.jp/entry/bps00401) |
| 10 | Phenazine biosynthesis | [bps00405](https://www.kegg.jp/entry/bps00405) |
| 11 | Cyanoamino acid metabolism | [bps00460](https://www.kegg.jp/entry/bps00460) |
| 12 | Streptomycin biosynthesis | [bps00521](https://www.kegg.jp/entry/bps00521) |
| 13 | Polyketide sugar unit biosynthesis | [bps00523](https://www.kegg.jp/entry/bps00523) |
| 14 | Acarbose and validamycin biosynthesis | [bps00525](https://www.kegg.jp/entry/bps00525) |
| 15 | Lipopolysaccharide biosynthesis | [bps00540](https://www.kegg.jp/entry/bps00540) |
| 16 | O-Antigen nucleotide sugar biosynthesis | [bps00541](https://www.kegg.jp/entry/bps00541) |
| 17 | Exopolysaccharide biosynthesis | [bps00543](https://www.kegg.jp/entry/bps00543) |
| 18 | Peptidoglycan biosynthesis | [bps00550](https://www.kegg.jp/entry/bps00550) |
| 19 | Teichoic acid biosynthesis | [bps00552](https://www.kegg.jp/entry/bps00552) |
| 20 | Dioxin degradation | [bps00621](https://www.kegg.jp/entry/bps00621) |
| 21 | Xylene degradation | [bps00622](https://www.kegg.jp/entry/bps00622) |
| 22 | Toluene degradation | [bps00623](https://www.kegg.jp/entry/bps00623) |
| 23 | Chloroalkane and chloroalkene degradation | [bps00625](https://www.kegg.jp/entry/bps00625) |
| 24 | Naphthalene degradation | [bps00626](https://www.kegg.jp/entry/bps00626) |
| 25 | Aminobenzoate degradation | [bps00627](https://www.kegg.jp/entry/bps00627) |
| 26 | Styrene degradation | [bps00643](https://www.kegg.jp/entry/bps00643) |
| 27 | C5-Branched dibasic acid metabolism | [bps00660](https://www.kegg.jp/entry/bps00660) |
| 28 | Methane metabolism | [bps00680](https://www.kegg.jp/entry/bps00680) |
| 29 | Carbon fixation in photosynthetic organisms | [bps00710](https://www.kegg.jp/entry/bps00710) |
| 30 | Atrazine degradation | [bps00791](https://www.kegg.jp/entry/bps00791) |
| 31 | Limonene and pinene degradation | [bps00903](https://www.kegg.jp/entry/bps00903) |
| 32 | Carotenoid biosynthesis | [bps00906](https://www.kegg.jp/entry/bps00906) |
| 33 | Sesquiterpenoid and triterpenoid biosynthesis | [bps00909](https://www.kegg.jp/entry/bps00909) |
| 34 | Caprolactam degradation | [bps00930](https://www.kegg.jp/entry/bps00930) |
| 35 | Degradation of flavonoids | [bps00946](https://www.kegg.jp/entry/bps00946) |
| 36 | Steroid degradation | [bps00984](https://www.kegg.jp/entry/bps00984) |
| 37 | Biosynthesis of various alkaloids | [bps00996](https://www.kegg.jp/entry/bps00996) |
| 38 | Biosynthesis of various other secondary metabolites | [bps00997](https://www.kegg.jp/entry/bps00997) |
| 39 | Biosynthesis of various antibiotics | [bps00998](https://www.kegg.jp/entry/bps00998) |
| 40 | Biosynthesis of various plant secondary metabolites | [bps00999](https://www.kegg.jp/entry/bps00999) |
| 41 | Biosynthesis of siderophore group nonribosomal peptides | [bps01053](https://www.kegg.jp/entry/bps01053) |
| 42 | Biosynthesis of secondary metabolites | [bps01110](https://www.kegg.jp/entry/bps01110) |
| 43 | Microbial metabolism in diverse environments | [bps01120](https://www.kegg.jp/entry/bps01120) |
| 44 | Degradation of aromatic compounds | [bps01220](https://www.kegg.jp/entry/bps01220) |
| 45 | beta-Lactam resistance | [bps01501](https://www.kegg.jp/entry/bps01501) |
| 46 | Vancomycin resistance | [bps01502](https://www.kegg.jp/entry/bps01502) |
| 47 | Cationic antimicrobial peptide (CAMP) resistance | [bps01503](https://www.kegg.jp/entry/bps01503) |
| 48 | Two-component system | [bps02020](https://www.kegg.jp/entry/bps02020) |
| 49 | Quorum sensing | [bps02024](https://www.kegg.jp/entry/bps02024) |
| 50 | Bacterial chemotaxis | [bps02030](https://www.kegg.jp/entry/bps02030) |
| 51 | Flagellar assembly | [bps02040](https://www.kegg.jp/entry/bps02040) |
| 52 | Phosphotransferase system (PTS) | [bps02060](https://www.kegg.jp/entry/bps02060) |
| 53 | Bacterial secretion system | [bps03070](https://www.kegg.jp/entry/bps03070) |

**Supplementary table 2:** Common pathways (KEGG) for *Burkholderia pseudomallei* K96243 (TaxID: 272560) compared to Homo sapiens (TaxID: 9606)

| **Sl No.** | **Common pathways (KEGG) for B. pseudomallei K96243** | **KEGG pathway ID** |
| --- | --- | --- |
| 1 | Glycolysis / Gluconeogenesis | [bps00010](https://www.kegg.jp/entry/bps00010) |
| 2 | Citrate cycle (TCA cycle) | [bps00020](https://www.kegg.jp/entry/bps00020) |
| 3 | Pentose phosphate pathway | [bps00030](https://www.kegg.jp/entry/bps00030) |
| 4 | Pentose and glucuronate interconversions | [bps00040](https://www.kegg.jp/entry/bps00040) |
| 5 | Fructose and mannose metabolism | [bps00051](https://www.kegg.jp/entry/bps00051) |
| 6 | Galactose metabolism | [bps00052](https://www.kegg.jp/entry/bps00052) |
| 7 | Ascorbate and aldarate metabolism | [bps00053](https://www.kegg.jp/entry/bps00053) |
| 8 | Fatty acid biosynthesis | [bps00061](https://www.kegg.jp/entry/bps00061) |
| 9 | Fatty acid degradation | [bps00071](https://www.kegg.jp/entry/bps00071) |
| 10 | Steroid biosynthesis | [bps00100](https://www.kegg.jp/entry/bps00100) |
| 11 | Ubiquinone and other terpenoid-quinone biosynthesis | [bps00130](https://www.kegg.jp/entry/bps00130) |
| 12 | Oxidative phosphorylation | [bps00190](https://www.kegg.jp/entry/bps00190) |
| 13 | Arginine biosynthesis | [bps00220](https://www.kegg.jp/entry/bps00220) |
| 14 | Purine metabolism | [bps00230](https://www.kegg.jp/entry/bps00230) |
| 15 | Pyrimidine metabolism | [bps00240](https://www.kegg.jp/entry/bps00240) |
| 16 | Alanine, aspartate and glutamate metabolism | [bps00250](https://www.kegg.jp/entry/bps00250) |
| 17 | Glycine, serine and threonine metabolism | [bps00260](https://www.kegg.jp/entry/bps00260) |
| 18 | Cysteine and methionine metabolism | [bps00270](https://www.kegg.jp/entry/bps00270) |
| 19 | Valine, leucine and isoleucine degradation | [bps00280](https://www.kegg.jp/entry/bps00280) |
| 20 | Valine, leucine and isoleucine biosynthesis | [bps00290](https://www.kegg.jp/entry/bps00290) |
| 21 | Lysine degradation | [bps00310](https://www.kegg.jp/entry/bps00310) |
| 22 | Arginine and proline metabolism | [bps00330](https://www.kegg.jp/entry/bps00330) |
| 23 | Histidine metabolism | [bps00340](https://www.kegg.jp/entry/bps00340) |
| 24 | Tyrosine metabolism | [bps00350](https://www.kegg.jp/entry/bps00350) |
| 25 | Phenylalanine metabolism | [bps00360](https://www.kegg.jp/entry/bps00360) |
| 26 | Tryptophan metabolism | [bps00380](https://www.kegg.jp/entry/bps00380) |
| 27 | Phenylalanine, tyrosine and tryptophan biosynthesis | [bps00400](https://www.kegg.jp/entry/bps00400) |
| 28 | beta-Alanine metabolism | [bps00410](https://www.kegg.jp/entry/bps00410) |
| 29 | Taurine and hypotaurine metabolism | [bps00430](https://www.kegg.jp/entry/bps00430) |
| 30 | Phosphonate and phosphinate metabolism | [bps00440](https://www.kegg.jp/entry/bps00440) |
| 31 | Selenocompound metabolism | [bps00450](https://www.kegg.jp/entry/bps00450) |
| 32 | D-Amino acid metabolism | [bps00470](https://www.kegg.jp/entry/bps00470) |
| 33 | Glutathione metabolism | [bps00480](https://www.kegg.jp/entry/bps00480) |
| 34 | Starch and sucrose metabolism | [bps00500](https://www.kegg.jp/entry/bps00500) |
| 35 | Other glycan degradation | [bps00511](https://www.kegg.jp/entry/bps00511) |
| 36 | Amino sugar and nucleotide sugar metabolism | [bps00520](https://www.kegg.jp/entry/bps00520) |
| 37 | Glycerolipid metabolism | [bps00561](https://www.kegg.jp/entry/bps00561) |
| 38 | Inositol phosphate metabolism | [bps00562](https://www.kegg.jp/entry/bps00562) |
| 39 | Glycerophospholipid metabolism | [bps00564](https://www.kegg.jp/entry/bps00564) |
| 40 | alpha-Linolenic acid metabolism | [bps00592](https://www.kegg.jp/entry/bps00592) |
| 41 | Sphingolipid metabolism | [bps00600](https://www.kegg.jp/entry/bps00600) |
| 42 | Pyruvate metabolism | [bps00620](https://www.kegg.jp/entry/bps00620) |
| 43 | Glyoxylate and dicarboxylate metabolism | [bps00630](https://www.kegg.jp/entry/bps00630) |
| 44 | Propanoate metabolism | [bps00640](https://www.kegg.jp/entry/bps00640) |
| 45 | Butanoate metabolism | [bps00650](https://www.kegg.jp/entry/bps00650) |
| 46 | One carbon pool by folate | [bps00670](https://www.kegg.jp/entry/bps00670) |
| 47 | Thiamine metabolism | [bps00730](https://www.kegg.jp/entry/bps00730) |
| 48 | Riboflavin metabolism | [bps00740](https://www.kegg.jp/entry/bps00740) |
| 49 | Vitamin B6 metabolism | [bps00750](https://www.kegg.jp/entry/bps00750) |
| 50 | Nicotinate and nicotinamide metabolism | [bps00760](https://www.kegg.jp/entry/bps00760) |
| 51 | Pantothenate and CoA biosynthesis | [bps00770](https://www.kegg.jp/entry/bps00770) |
| 52 | Biotin metabolism | [bps00780](https://www.kegg.jp/entry/bps00780) |
| 53 | Lipoic acid metabolism | [bps00785](https://www.kegg.jp/entry/bps00785) |
| 54 | Folate biosynthesis | [bps00790](https://www.kegg.jp/entry/bps00790) |
| 55 | Porphyrin metabolism | [bps00860](https://www.kegg.jp/entry/bps00860) |
| 56 | Terpenoid backbone biosynthesis | [bps00900](https://www.kegg.jp/entry/bps00900) |
| 57 | Nitrogen metabolism | [bps00910](https://www.kegg.jp/entry/bps00910) |
| 58 | Sulfur metabolism | [bps00920](https://www.kegg.jp/entry/bps00920) |
| 59 | Aminoacyl-tRNA biosynthesis | [bps00970](https://www.kegg.jp/entry/bps00970) |
| 60 | Biosynthesis of unsaturated fatty acids | [bps01040](https://www.kegg.jp/entry/bps01040) |
| 61 | Metabolic pathways | [bps01100](https://www.kegg.jp/entry/bps01100) |
| 62 | Carbon metabolism | [bps01200](https://www.kegg.jp/entry/bps01200) |
| 63 | 2-Oxocarboxylic acid metabolism | [bps01210](https://www.kegg.jp/entry/bps01210) |
| 64 | Fatty acid metabolism | [bps01212](https://www.kegg.jp/entry/bps01212) |
| 65 | Biosynthesis of amino acids | [bps01230](https://www.kegg.jp/entry/bps01230) |
| 66 | Nucleotide metabolism | [bps01232](https://www.kegg.jp/entry/bps01232) |
| 67 | Biosynthesis of cofactors | [bps01240](https://www.kegg.jp/entry/bps01240) |
| 68 | Biosynthesis of nucleotide sugars | [bps01250](https://www.kegg.jp/entry/bps01250) |
| 69 | ABC transporters | [bps02010](https://www.kegg.jp/entry/bps02010) |
| 70 | Ribosome | [bps03010](https://www.kegg.jp/entry/bps03010) |
| 71 | RNA degradation | [bps03018](https://www.kegg.jp/entry/bps03018) |
| 72 | RNA polymerase | [bps03020](https://www.kegg.jp/entry/bps03020) |
| 73 | DNA replication | [bps03030](https://www.kegg.jp/entry/bps03030) |
| 74 | Protein export | [bps03060](https://www.kegg.jp/entry/bps03060) |
| 75 | Viral life cycle - HIV-1 | [bps03250](https://www.kegg.jp/entry/bps03250) |
| 76 | Base excision repair | [bps03410](https://www.kegg.jp/entry/bps03410) |
| 77 | Nucleotide excision repair | [bps03420](https://www.kegg.jp/entry/bps03420) |
| 78 | Mismatch repair | [bps03430](https://www.kegg.jp/entry/bps03430) |
| 79 | Homologous recombination | [bps03440](https://www.kegg.jp/entry/bps03440) |
| 80 | Non-homologous end-joining | [bps03450](https://www.kegg.jp/entry/bps03450) |
| 81 | Sulfur relay system | [bps04122](https://www.kegg.jp/entry/bps04122) |

**Supplementary table 3:** Complete list of essential non-homologous genes in *Burkholderia pseudomallei* K96243

| **Sl No.** | **UniProt ID** | **Protein** | **Sub-cellular localization** | **TMHMM** | **Drugbank ID** |
| --- | --- | --- | --- | --- | --- |
| 1 | [Q63YW7](https://www.uniprot.org/uniprotkb/Q63YW7/entry) | DNA gyrase subunit B (gyrB) | Cytoplasmic | No | DB00817 DB04395 DB03966 DB05488 DB01044 DB01051 |
| 2 | [Q63YW6](https://www.uniprot.org/uniprot/Q63YW6) | Beta sliding clamp (dnaN) | Cytoplasmic | No | No |
| 3 | [Q63YW5](https://www.uniprot.org/uniprot/Q63YW5) | Chromosomal replication initiator protein DnaA (dnaA) | Cytoplasmic | No | No |
| 4 | [Q63YW1](https://www.uniprot.org/uniprot/Q63YW1) | Membrane protein insertase YidC (yidC) | Inner Membrane | Yes | No |
| 5 | [Q63YV0](https://www.uniprot.org/uniprot/Q63YV0) | Uncharacterized protein | Cytoplasmic | No | No |
| 6 | [Q63YU9](https://www.uniprot.org/uniprot/Q63YU9) | Transposase protein | Cytoplasmic | No | No |
| 7 | [Q63YR0](https://www.uniprot.org/uniprot/Q63YR0) | Response regulator protein | Cytoplasmic | No | No |
| 8 | [Q63YK1](https://www.uniprot.org/uniprot/Q63YK1) | Cell shape-determining protein MreB (mreB) | Cytoplasmic | No | No |
| 9 | [Q63YJ9](https://www.uniprot.org/uniprot/Q63YJ9) | Glutamyl-tRNA(Gln) amidotransferase subunit A (gatA) | Periplasmic | No | No |
| 10 | [Q63YI7](https://www.uniprot.org/uniprot/Q63YI7) | Acetylglutamate kinase (argB) | Cytoplasmic | No | No |
| 11 | [Q63YH9](https://www.uniprot.org/uniprot/Q63YH9) | Tyrosine recombinase XerC (xerC) | Cytoplasmic | No | No |
| 12 | [Q63YE7](https://www.uniprot.org/uniprot/Q63YE7) | ferredoxin--NADP(+) reductase (fpr) | Cytoplasmic | No | DB03147 |
| 13 | [Q63YC3](https://www.uniprot.org/uniprot/Q63YC3) | Multifunctional CCA protein (cca) | Cytoplasmic | No | No |
| 14 | [Q63Y76](https://www.uniprot.org/uniprot/Q63Y76) | Glutamine--fructose-6-phosphate aminotransferase [isomerizing] (glmS1) | Cytoplasmic | No | DB00130 |
| 15 | [Q63Y75](https://www.uniprot.org/uniprot/Q63Y75) | Bifunctional protein GlmU (glmU) | Cytoplasmic | No | No |
| 16 | [Q63Y71](https://www.uniprot.org/uniprot/Q63Y71) | SAM-dependent methyltransferase | Cytoplasmic | No | No |
| 17 | [Q63Y06](https://www.uniprot.org/uniprot/Q63Y06) | Arginine--tRNA ligase (argS) | Cytoplasmic | No | No |
| 18 | [Q63XZ2](https://www.uniprot.org/uniprot/Q63XZ2) | Type III pantothenate kinase (coaX) | Cytoplasmic | No | No |
| 19 | [Q63XZ1](https://www.uniprot.org/uniprot/Q63XZ1) | biotin--[biotin carboxyl-carrier protein] ligase | Cytoplasmic | No | No |
| 20 | [Q63XM5](https://www.uniprot.org/uniprot/Q63XM5) | Signal recognition particle receptor FtsY (ftsY) | Cytoplasmic | No | DB04272 |
| 21 | [Q63XL8](https://www.uniprot.org/uniprot/Q63XL8) | Ribose-phosphate pyrophosphokinase (prs) | Cytoplasmic | No | DB11638 |
| 22 | [Q63XL7](https://www.uniprot.org/uniprot/Q63XL7) | 4-diphosphocytidyl-2-C-methyl-D-erythritol kinase (ispE) | Cytoplasmic | No | No |
| 23 | [Q63XL0](https://www.uniprot.org/uniprot/Q63XL0) | HPr kinase/phosphorylase (hprK) | Cytoplasmic | No | No |
| 24 | [Q63XK](https://www.uniprot.org/uniprot/Q63XK6)6 | ABC transporter system, ATP-binding protein | Cytoplasmic | No | DB05154 |
| 25 | [Q63XK](https://www.uniprot.org/uniprot/Q63XK5)5 | Lipopolysaccharide export system protein LptA (lptA) | Periplasmic | No | No |
| 26 | [Q63XJ3](https://www.uniprot.org/uniprot/Q63XJ3) | Single-stranded DNA-binding protein (ssb) | Extracellular | No | No |
| 27 | [Q63XF3](https://www.uniprot.org/uniprot/Q63XF3) | Membrane protein | Outer Membrane | Yes | No |
| 28 | [Q63XE](https://www.uniprot.org/uniprot/Q63XE8)8 | DUF4760 domain-containing protein | Cytoplasmic | No | No |
| 29 | [Q63XE3](https://www.uniprot.org/uniprot/Q63XE3) | Uncharacterized protein | Cytoplasmic | No | No |
| 30 | [Q63XD4](https://www.uniprot.org/uniprot/Q63XD4) | Membrane protein | Cytoplasmic | Yes | No |
| 31 | [Q63XD3](https://www.uniprot.org/uniprot/Q63XD3) | Polysaccharide biosynthesis protein | Inner Membrane | Yes | No |
| 32 | [Q63XD2](https://www.uniprot.org/uniprot/Q63XD2) | Glycosyl transferase | Cytoplasmic | No | No |
| 33 | [Q63X97](https://www.uniprot.org/uniprot/Q63X97) | Probable protein kinase UbiB (ubiB) | Inner Membrane | Yes | No |
| 34 | [Q63X94](https://www.uniprot.org/uniprot/Q63X94) | Membrane protein | Inner Membrane | Yes | No |
| 35 | [Q63X93](https://www.uniprot.org/uniprot/Q63X93) | Aspartate--tRNA(Asp/Asn) ligase (aspS) | Cytoplasmic | No | DB00128 |
| 36 | [Q63X79](https://www.uniprot.org/uniprot/Q63X79) | LPS-assembly protein LptD (lptD) | Outer Membrane | No | No |
| 37 | [Q63X77](https://www.uniprot.org/uniprot/Q63X77) | 4-hydroxythreonine-4-phosphate dehydrogenase (pdxA) | Cytoplasmic | No | No |
| 38 | [Q63X70](https://www.uniprot.org/uniprot/Q63X70) | Glycine--tRNA ligase beta subunit (glyS) | Cytoplasmic | No | No |
| 39 | [Q63X65](https://www.uniprot.org/uniprot/Q63X65) | Endoribonuclease YbeY (ybeY) | Cytoplasmic | No | No |
| 40 | [Q63WV0](https://www.uniprot.org/uniprot/Q63WV0) | ADP-heptose--LPS heptosyltransferase II (waaF) | Cytoplasmic | No | No |
| 41 | [Q63WU0](https://www.uniprot.org/uniprot/Q63WU0) | N5-carboxyaminoimidazole ribonucleotide synthase (purK) | Cytoplasmic | No | No |
| 42 | [Q63WT9](https://www.uniprot.org/uniprot/Q63WT9) | Threonylcarbamoyl-AMP synthase | Cytoplasmic | No | No |
| 43 | [Q63WN2](https://www.uniprot.org/uniprot/Q63WN2) | N-acetylmuramoyl-L-alanine amidase | Extracellular | No | No |
| 44 | [Q63WN1](https://www.uniprot.org/uniprot/Q63WN1) | tRNA threonylcarbamoyladenosine biosynthesis protein TsaE | Cytoplasmic | No | No |
| 45 | [Q63WM5](https://www.uniprot.org/uniprot/Q63WM5) | Glycerol-3-phosphate acyltransferase (plsY) | Inner Membrane | Yes | No |
| 46 | [Q63WM3](https://www.uniprot.org/uniprot/Q63WM3) | UDP-N-acetylenolpyruvoylglucosamine reductase (murB) | Cytoplasmic | No | DB03147 DB07296 |
| 47 | [Q63WL9](https://www.uniprot.org/uniprot/Q63WL9) | Probable lipid II flippase MurJ (murJ) | Extracellular | Yes | No |
| 48 | [Q63WL6](https://www.uniprot.org/uniprot/Q63WL6) | Adenylate kinase (adk) | Cytoplasmic | No | No |
| 49 | [Q63WL3](https://www.uniprot.org/uniprot/Q63WL3) | Tetraacyldisaccharide 4'-kinase (lpxK) | Cytoplasmic | Yes | No |
| 50 | [Q63WL1](https://www.uniprot.org/uniprot/Q63WL1) | Superoxide dismutase (sodB) | Extracellular | No | No |
| 51 | [Q63WJ4](https://www.uniprot.org/uniprot/Q63WJ4) | Isocitrate dehydrogenase [NADP] (icd) | Cytoplasmic | No | DB02159 DB04272 DB04349 DB04530 DB00157 DB09130 DB06757 DB09092 |
| 52 | [Q63WI5](https://www.uniprot.org/uniprot/Q63WI5) | Coenzyme A biosynthesis bifunctional protein CoaBC (dfp) | Cytoplasmic | No | DB03738 DB03247 |
| 53 | [Q63WI4](https://www.uniprot.org/uniprot/Q63WI4) | Lipoprotein signal peptidase (lspA) | Inner Membrane | Yes | No |
| 54 | [Q63WI3](https://www.uniprot.org/uniprot/Q63WI3) | Isoleucine--tRNA ligase 1 (ileS1) | Cytoplasmic | No | DB00410 DB11638 |
| 55 | [Q63WC5](https://www.uniprot.org/uniprot/Q63WC5) | Permease protein | Inner Membrane | Yes | No |
| 56 | [Q63WC4](https://www.uniprot.org/uniprot/Q63WC4) | Lipopolysaccharide export system permease protein LptF | Inner Membrane | Yes | No |
| 57 | [Q63WB9](https://www.uniprot.org/uniprot/Q63WB9) | Dihydroxy-acid dehydratase (ilvD) | Cytoplasmic | No | No |
| 58 | [Q63W96](https://www.uniprot.org/uniprot/Q63W96) | Segregation and condensation protein A | Cytoplasmic | No | No |
| 59 | [Q63W86](https://www.uniprot.org/uniprot/Q63W86) | dCTP deaminase (dcd) | Cytoplasmic | No | No |
| 60 | [Q63W6](https://www.uniprot.org/uniprot/Q63W64)4 | Glutamine-dependent NAD(+) synthetase (nadE) | Cytoplasmic | No | No |
| 61 | [Q63W24](https://www.uniprot.org/uniprot/Q63W24) | DNA topoisomerase 4 subunit B (parE) | Cytoplasmic | No | DB00537 DB01044 |
| 62 | [Q63VX8](https://www.uniprot.org/uniprot/Q63VX8) | DNA polymerase III subunit alpha (dnaE) | Cytoplasmic | No | No |
| 63 | [Q63VX](https://www.uniprot.org/uniprot/Q63VX7)7 | ATP-dependent lipid A-core flippase (msbA) | Inner Membrane | Yes | DB05154 |
| 64 | [Q63VT2](https://www.uniprot.org/uniprot/Q63VT2) | Probable nicotinate-nucleotide adenylyltransferase (nadD) | Cytoplasmic | No | DB04272 |
| 65 | [Q63VP6](https://www.uniprot.org/uniprot/Q63VP6) | Ketol-acid reductoisomerase (NADP(+)) (ilvC) | Cytoplasmic | No | No |
| 66 | [Q63VP5](https://www.uniprot.org/uniprot/Q63VP5) | Phosphatidylserine decarboxylase proenzyme (psd) | Inner Membrane | Yes | No |
| 67 | [Q63VN7](https://www.uniprot.org/uniprot/Q63VN7) | Polyribonucleotide nucleotidyltransferase (pnp) | Cytoplasmic | No | No |
| 68 | [Q63VM7](https://www.uniprot.org/uniprot/Q63VM7) | NADH-quinone oxidoreductase | Cytoplasmic | No | No |
| 69 | [Q63VM4](https://www.uniprot.org/uniprot/Q63VM4) | NADH-quinone oxidoreductase subunit J (nuoJ) | Inner Membrane | Yes | No |
| 70 | [Q63VM1](https://www.uniprot.org/uniprot/Q63VM1) | NADH dehydrogenase I chain M (nuoM) | Inner Membrane | Yes | No |
| 71 | [Q63VM0](https://www.uniprot.org/uniprot/Q63VM0) | NADH-quinone oxidoreductase subunit N (nuoN) | Inner Membrane | Yes | No |
| 72 | [Q63VG3](https://www.uniprot.org/uniprot/Q63VG3) | RecBCD enzyme subunit RecC (recC) | Cytoplasmic | No | No |
| 73 | [Q63VA0](https://www.uniprot.org/uniprot/Q63VA0) | DNA polymerase III subunit epsilon (dnaQ) | Cytoplasmic | No | No |
| 74 | [Q63V99](https://www.uniprot.org/uniprot/Q63V99) | Ribonuclease H (rnhA) | Extracellular | No | No |
| 75 | [Q63V86](https://www.uniprot.org/uniprot/Q63V86) | Ribosomal RNA large subunit methyltransferase E (rlmE) | Cytoplasmic | No | No |
| 76 | [Q63V84](https://www.uniprot.org/uniprot/Q63V84) | Dihydropteroate synthase (folP) | Cytoplasmic | No | DB00576 DB01298 DB00634 DB00259 DB01015 DB01581 DB01582 DB06729 DB00250 DB00664 DB01299 DB01145 DB00359 DB08798 DB06147 |
| 77 | [Q63V83](https://www.uniprot.org/uniprot/Q63V83) | Phosphoglucosamine mutase (glmM) | Cytoplasmic | No | No |
| 78 | [Q63V54](https://www.uniprot.org/uniprot/Q63V54) | DUF4123 domain-containing protein | Cytoplasmic | No | No |
| 79 | [Q63V40](https://www.uniprot.org/uniprot/Q63V40) | ATP-dependent Clp protease ATP-binding subunit ClpX (clpX) | Cytoplasmic | No | DB09275 |
| 80 | [Q63V10](https://www.uniprot.org/uniprot/Q63V10) | Uncharacterized protein | Cytoplasmic | No | No |
| 81 | [Q63V08](https://www.uniprot.org/uniprot/Q63V08) | Thymidylate kinase (tmk) | Cytoplasmic | No | No |
| 82 | [Q63UY2](https://www.uniprot.org/uniprot/Q63UY2) | Replicative DNA helicase (dnaB) | Cytoplasmic | No | No |
| 83 | [Q63UX6](https://www.uniprot.org/uniprot/Q63UX6) | NodB homology domain-containing protein | Cytoplasmic | No | No |
| 84 | [Q63UX5](https://www.uniprot.org/uniprot/Q63UX5) | Polymixin resistance glucosyl transferase | Cytoplasmic | Yes | No |
| 85 | [Q63UX4](https://www.uniprot.org/uniprot/Q63UX4) | LPS biosynthesis-related protein | Cytoplasmic | No | No |
| 86 | [Q63UX2](https://www.uniprot.org/uniprot/Q63UX2) | Membrane protein | Inner Membrane | Yes | No |
| 87 | [Q63UW9](https://www.uniprot.org/uniprot/Q63UW9) | Homoserine dehydrogenase (hom) | Cytoplasmic | No | No |
| 88 | [Q63UW8](https://www.uniprot.org/uniprot/Q63UW8) | Threonine synthase (thrC) | Cytoplasmic | No | No |
| 89 | [Q63UV1](https://www.uniprot.org/uniprot/Q63UV1) | Transcription termination factor Rho (rho) | Cytoplasmic | No | No |
| 90 | [Q63UU8](https://www.uniprot.org/uniprot/Q63UU8) | DNA polymerase III subunit gamma (dnaX) | Cytoplasmic | No | No |
| 91 | [Q63UU2](https://www.uniprot.org/uniprot/Q63UU2) | Subfamily M23B unassigned peptidase | Extracellular | No | No |
| 92 | [Q63UT6](https://www.uniprot.org/uniprot/Q63UT6) | Nucleoside diphosphate kinase (ndk) | Extracellular | No | DB04315 DB00709 DB09299 DB01262 DB00787 |
| 93 | [Q63UT3](https://www.uniprot.org/uniprot/Q63UT3) | 4-hydroxy-3-methylbut-2-en-1-yl diphosphate synthase (flavodoxin) (ispG) | Cytoplasmic | No | No |
| 94 | [Q63UT](https://www.uniprot.org/uniprot/Q63UT2)2 | Histidine--tRNA ligase (hisS) | Cytoplasmic | No | No |
| 95 | [Q63UT0](https://www.uniprot.org/uniprot/Q63UT0) | Outer membrane protein assembly factor BamB (bamB) | Periplasmic | No | No |
| 96 | [Q63UK](https://www.uniprot.org/uniprot/Q63UK2)2 | Transport-related, membrane protein | Inner Membrane | Yes | No |
| 97 | [Q63UK1](https://www.uniprot.org/uniprot/Q63UK1) | Membrane linked regulatory protein | Inner Membrane | Yes | No |
| 98 | [Q63UK](https://www.uniprot.org/uniprot/Q63UK0)0 | Cytochrome c-related lipoprotein | Periplasmic | No | No |
| 99 | [Q63UJ7](https://www.uniprot.org/uniprot/Q63UJ7) | Lipoprotein | Periplasmic | No | No |
| 100 | [Q63UJ6](https://www.uniprot.org/uniprot/Q63UJ6) | Membrane protein | Inner Membrane | Yes | No |
| 101 | [Q63UJ](https://www.uniprot.org/uniprot/Q63UJ5)5 | Copper-related ABC transport system, ATP-binding protein | Cytoplasmic | No | No |
| 102 | [Q63UJ4](https://www.uniprot.org/uniprot/Q63UJ4) | Copper-binding periplasmic protein | Periplasmic | No | No |
| 103 | [Q63UJ2](https://www.uniprot.org/uniprot/Q63UJ2) | FAD:protein FMN transferase | Cytoplasmic | Yes | No |
| 104 | [Q63UJ0](https://www.uniprot.org/uniprot/Q63UJ0) | Transport permease protein | Inner Membrane | Yes | No |
| 105 | [Q63UI9](https://www.uniprot.org/uniprot/Q63UI9) | Methyl-accepting chemotaxis transmembrane protein | Inner Membrane | Yes | No |
| 106 | [Q63UI8](https://www.uniprot.org/uniprot/Q63UI8) | Uncharacterized protein | Cytoplasmic | No | No |
| 107 | [Q63UI5](https://www.uniprot.org/uniprot/Q63UI5) | Iron-sulphur protein | Cytoplasmic | No | No |
| 108 | [Q63UI](https://www.uniprot.org/uniprot/Q63UI4)4 | Transport related membrane protein | Inner Membrane | Yes | No |
| 109 | [Q63UI3](https://www.uniprot.org/uniprot/Q63UI3) | Hydroxylase | Cytoplasmic | No | No |
| 110 | [Q63UI2](https://www.uniprot.org/uniprot/Q63UI2) | Iron-containing redox enzyme family protein | Cytoplasmic | No | No |
| 111 | [Q63UH5](https://www.uniprot.org/uniprot/Q63UH5) | Fimbrial assembly chaperone | Periplasmic | No | No |
| 112 | [Q63UH4](https://www.uniprot.org/uniprot/Q63UH4) | Fimbrial usher protein | Outer Membrane | No | No |
| 113 | [Q63UH3](https://www.uniprot.org/uniprot/Q63UH3) | Exported fimbria-related protein | Extracellular | Yes | No |
| 114 | [Q63UH1](https://www.uniprot.org/uniprot/Q63UH1) | Outer membrane protein | Extracellular | No | No |
| 115 | [Q63UG3](https://www.uniprot.org/uniprot/Q63UG3) | Amino acid synthesis family protein | Cytoplasmic | No | No |
| 116 | [Q63UG1](https://www.uniprot.org/uniprot/Q63UG1) | Oxygenase | Cytoplasmic | No | No |
| 117 | [Q63UF7](https://www.uniprot.org/uniprot/Q63UF7) | ABC transport system, substrate-binding protein | Periplasmic | No | No |
| 118 | [Q63UF](https://www.uniprot.org/uniprot/Q63UF6)6 | ABC transport system, permease protein | Inner Membrane | Yes | No |
| 119 | [Q63UF5](https://www.uniprot.org/uniprot/Q63UF5) | ABC transport system, permease protein | Inner Membrane | Yes | No |
| 120 | [Q63UF4](https://www.uniprot.org/uniprot/Q63UF4) | ABC transport system, ATP-binding protein | Inner Membrane | No | DB05154 |
| 121 | [Q63UF3](https://www.uniprot.org/uniprot/Q63UF3) | GntR-family regulatory protein | Cytoplasmic | No | No |
| 122 | [Q63UF2](https://www.uniprot.org/uniprot/Q63UF2) | Succinate-semialdehyde dehydrogenase [NADP+] (gabD) | Cytoplasmic | No | DB00534 DB00157 DB00139 DB00313 DB09072 |
| 123 | [Q63UF1](https://www.uniprot.org/uniprot/Q63UF1) | Outer membrane porin protein | Outer Membrane | No | No |
| 124 | [Q63UE9](https://www.uniprot.org/uniprot/Q63UE9) | Integrase catalytic domain-containing protein | Cytoplasmic | No | No |
| 125 | [Q63UE6](https://www.uniprot.org/uniprot/Q63UE6) | Exported protein | Outer Membrane | Yes | No |
| 126 | [Q63UE](https://www.uniprot.org/uniprot/Q63UE5)5 | Outer membrane protein | Outer Membrane | No | No |
| 127 | [Q63UE4](https://www.uniprot.org/uniprot/Q63UE4) | Hemolysin-related protein | Extracellular | No | No |
| 128 | [Q63UE3](https://www.uniprot.org/uniprot/Q63UE3) | Fe2OG dioxygenase domain-containing protein | Cytoplasmic | No | No |
| 129 | [Q63UE2](https://www.uniprot.org/uniprot/Q63UE2) | Sulfotransferase | Cytoplasmic | No | No |
| 130 | [Q63UE1](https://www.uniprot.org/uniprot/Q63UE1) | Toxin transport-related membrane protein | Inner Membrane | Yes | No |
| 131 | [Q63UE0](https://www.uniprot.org/uniprot/Q63UE0) | Membrane fusion protein (MFP) family protein | Inner Membrane | Yes | No |
| 132 | [Q63UD9](https://www.uniprot.org/uniprot/Q63UD9) | Methyltransferase type 11 domain-containing protein | Cytoplasmic | No | No |
| 133 | [Q63UD8](https://www.uniprot.org/uniprot/Q63UD8) | Tetratricopeptide repeat protein | Cytoplasmic | No | No |
| 134 | [Q63UD](https://www.uniprot.org/uniprot/Q63UD5)5 | Transposase | Cytoplasmic | No | No |
| 135 | [Q63UD3](https://www.uniprot.org/uniprot/Q63UD3) | DUF3443 domain-containing protein | Extracellular | No | No |
| 136 | [Q63UD2](https://www.uniprot.org/uniprot/Q63UD2) | Outer membrane porin protein | Outer Membrane | No | No |
| 137 | [Q63UD1](https://www.uniprot.org/uniprot/Q63UD1) | Transposase | Cytoplasmic | No | No |
| 138 | [Q63UD](https://www.uniprot.org/uniprot/Q63UD0)0 | Exported histidine ammonia-lyase | Cytoplasmic | No | No |
| 139 | [Q63UC9](https://www.uniprot.org/uniprot/Q63UC9) | Transport-related, integral membrane protein | Inner Membrane | Yes | No |
| 140 | [Q63UC](https://www.uniprot.org/uniprot/Q63UC8)8 | Urocanate hydratase | Cytoplasmic | No | No |
| 141 | [Q63UC7](https://www.uniprot.org/uniprot/Q63UC7) | DUF917 domain-containing protein | Cytoplasmic | No | No |
| 142 | [Q63UC5](https://www.uniprot.org/uniprot/Q63UC5) | Allantoinase | Cytoplasmic | No | No |
| 143 | [Q63UC4](https://www.uniprot.org/uniprot/Q63UC4) | Family M20 unassigned peptidase | Extracellular | No | No |
| 144 | [Q63UC0](https://www.uniprot.org/uniprot/Q63UC0) | Membrane protein | Inner Membrane | Yes | No |
| 145 | [Q63UB9](https://www.uniprot.org/uniprot/Q63UB9) | Putative DNA-binding domain-containing protein | Cytoplasmic | No | No |
| 146 | [Q63UB8](https://www.uniprot.org/uniprot/Q63UB8) | UPF0276 protein BPSL1691 | Cytoplasmic | No | No |
| 147 | [Q63UB1](https://www.uniprot.org/uniprot/Q63UB1) | Uncharacterized protein | Cytoplasmic | No | No |
| 148 | [Q63UB0](https://www.uniprot.org/uniprot/Q63UB0) | Membrane protein | Extracellular | Yes | No |
| 149 | [Q63UA7](https://www.uniprot.org/uniprot/Q63UA7) | Exported protein | Extracellular | No | No |
| 150 | [Q63UA0](https://www.uniprot.org/uniprot/Q63UA0) | Cysteine synthase | Cytoplasmic | No | No |
| 151 | [Q63U99](https://www.uniprot.org/uniprot/Q63U99) | Uncharacterized protein | Cytoplasmic | No | No |
| 152 | [Q63U98](https://www.uniprot.org/uniprot/Q63U98) | Membrane protein | Inner Membrane | Yes | No |
| 153 | [Q63U96](https://www.uniprot.org/uniprot/Q63U96) | Argininosuccinate lyase | Cytoplasmic | No | No |
| 154 | [Q63U95](https://www.uniprot.org/uniprot/Q63U95) | Argininosuccinate synthase | Cytoplasmic | No | DB00125 DB00128 DB00155 DB00171 |
| 155 | [Q63U94](https://www.uniprot.org/uniprot/Q63U94) | Formyl transferase | Cytoplasmic | No | No |
| 156 | [Q63U93](https://www.uniprot.org/uniprot/Q63U93) | Uncharacterized protein | Cytoplasmic | No | No |
| 157 | [Q63U92](https://www.uniprot.org/uniprot/Q63U92) | Aminotransferase | Cytoplasmic | No | No |
| 158 | [Q63U89](https://www.uniprot.org/uniprot/Q63U89) | Non-ribosomal peptide synthase (Thioesterase domain) | Cytoplasmic | No | No |
| 159 | [Q63U88](https://www.uniprot.org/uniprot/Q63U88) | Exported porin | Extracellular | No | No |
| 160 | [Q63U87](https://www.uniprot.org/uniprot/Q63U87) | AraC-family transcriptional regulator | Cytoplasmic | No | No |
| 161 | [Q63U86](https://www.uniprot.org/uniprot/Q63U86) | Transmembrane protein | Inner Membrane | Yes | No |
| 162 | [Q63U85](https://www.uniprot.org/uniprot/Q63U85) | Chemotaxis protein CheW2 (cheW2) | Cytoplasmic | No | No |
| 163 | [Q63U84](https://www.uniprot.org/uniprot/Q63U84) | Methyl-accepting chemotaxis citrate transducer | Inner Membrane | Yes | No |
| 164 | [Q63U7](https://www.uniprot.org/uniprot/Q63U74)4 | Arginine/ornithine antiporter (arcD) | Inner Membrane | Yes | No |
| 165 | [Q63U73](https://www.uniprot.org/uniprot/Q63U73) | Arginine deiminase (arcA) | Cytoplasmic | No | No |
| 166 | [Q63U66](https://www.uniprot.org/uniprot/Q63U66) | MarR-family transcriptional regulator | Cytoplasmic | No | No |
| 167 | [Q63U65](https://www.uniprot.org/uniprot/Q63U65) | Amino-acid transport-related exported protein | Periplasmic | No | No |
| 168 | [Q63U37](https://www.uniprot.org/uniprot/Q63U37) | Siderophore biosynthesis related ABC transport protein | Inner Membrane | Yes | No |
| 169 | [Q63U33](https://www.uniprot.org/uniprot/Q63U33) | Iron transport-related membrane protein | Inner Membrane | Yes | No |
| 170 | [Q63U32](https://www.uniprot.org/uniprot/Q63U32) | Iron transport-related ATP-bidning protein | Cytoplasmic | No | No |
| 171 | [Q63U25](https://www.uniprot.org/uniprot/Q63U25) | Uncharacterized protein | Cytoplasmic | No | No |
| 172 | [Q63U24](https://www.uniprot.org/uniprot/Q63U24) | Sugar-binding exported protein | Periplasmic | No | No |
| 173 | [Q63U23](https://www.uniprot.org/uniprot/Q63U23) | AraC-family transcriptional regulator | Cytoplasmic | No | No |
| 174 | [Q63U14](https://www.uniprot.org/uniprot/Q63U14) | Efflux pump membrane transporter (amrB) | Inner Membrane | Yes | No |
| 175 | [Q63U13](https://www.uniprot.org/uniprot/Q63U13) | Multidrug efflux system putative membrane fusion protein (amrA) | Inner Membrane | No | No |
| 176 | [Q63U07](https://www.uniprot.org/uniprot/Q63U07) | Membrane protein | Inner Membrane | Yes | No |
| 177 | [Q63U06](https://www.uniprot.org/uniprot/Q63U06) | Membrane protein | Inner Membrane | Yes | No |
| 178 | [Q63U04](https://www.uniprot.org/uniprot/Q63U04) | Fimbriae-related membrane protein | Inner Membrane | Yes | No |
| 179 | [Q63U03](https://www.uniprot.org/uniprot/Q63U03) | Membrane protein | Inner Membrane | Yes | No |
| 180 | [Q63U02](https://www.uniprot.org/uniprot/Q63U02) | Fimbriae assembly-related protein | Cytoplasmic | No | No |
| 181 | [Q63U01](https://www.uniprot.org/uniprot/Q63U01) | Fimbriae assembly-related protein | Cytoplasmic | No | No |
| 182 | [Q63TZ9](https://www.uniprot.org/uniprot/Q63TZ9) | Fimbriae assembly-related protein | Outer Membrane | No | No |
| 183 | [Q63TZ8](https://www.uniprot.org/uniprot/Q63TZ8) | Fimbriae assembly-related protein | Periplasmic | Yes | No |
| 184 | [Q63TZ7](https://www.uniprot.org/uniprot/Q63TZ7) | Fimbriae assembly-related protein | Inner Membrane | Yes | No |
| 185 | [Q63TZ5](https://www.uniprot.org/uniprot/Q63TZ5) | Membrane protein | Inner Membrane | Yes | No |
| 186 | [Q63TZ4](https://www.uniprot.org/uniprot/Q63TZ4) | ABC transport system, ATP-binding protein | Cytoplasmic | No | DB05154 |
| 187 | [Q63TU0](https://www.uniprot.org/uniprot/Q63TU0) | PA domain-containing protein | Cytoplasmic | No | No |
| 188 | [Q63TQ8](https://www.uniprot.org/uniprot/Q63TQ8) | Dihydrolipoyllysine-residue succinyltransferase component of 2-oxoglutarate dehydrogenase complex (sucB) | Periplasmic | No | No |
| 189 | [Q63TQ7](https://www.uniprot.org/uniprot/Q63TQ7) | oxoglutarate dehydrogenase (succinyl-transferring) (sucA) | Cytoplasmic | No | DB00157 DB00313 DB09092 |
| 190 | [Q63TQ6](https://www.uniprot.org/uniprot/Q63TQ6) | Transposase | Cytoplasmic | No | No |
| 191 | [Q63TP8](https://www.uniprot.org/uniprot/Q63TP8) | Translation initiation factor IF-2 (infB) | Cytoplasmic | No | No |
| 192 | [Q63R56](https://www.uniprot.org/uniprot/Q63R56) | Transposase (tnpA) | Cytoplasmic | No | No |
| 193 | [Q63TM7](https://www.uniprot.org/uniprot/Q63TM7) | Phenylalanine--tRNA ligase beta subunit (pheT) | Cytoplasmic | No | DB00120 |
| 194 | [Q63TM6](https://www.uniprot.org/uniprot/Q63TM6) | Phenylalanine--tRNA ligase alpha subunit (pheS) | Cytoplasmic | No | DB00120 |
| 195 | [Q63TM2](https://www.uniprot.org/uniprot/Q63TM2) | Threonine--tRNA ligase (thrS) | Cytoplasmic | No | DB00156 DB11638 |
| 196 | [Q63TI8](https://www.uniprot.org/uniprot/Q63TI8) | Valine--tRNA ligase (valS) | Cytoplasmic | No | DB00161 |
| 197 | [Q63TH](https://www.uniprot.org/uniprot/Q63TH5)5 | ABC transport system, ATP-binding protein | Cytoplasmic | No | DB05154 |
| 198 | [Q63TG1](https://www.uniprot.org/uniprot/Q63TG1) | Membrane protein | Inner Membrane | Yes | No |
| 199 | [Q63TF9](https://www.uniprot.org/uniprot/Q63TF9) | Alanine--tRNA ligase (alaS) | Cytoplasmic | No | No |
| 200 | [Q63TF6](https://www.uniprot.org/uniprot/Q63TF6) | Glutamine--tRNA ligase (glnS) | Cytoplasmic | No | DB00130 |
| 201 | [Q63TE8](https://www.uniprot.org/uniprot/Q63TE8) | HlyD family secretion protein | Outer Membrane | Yes | No |
| 202 | [Q63TE7](https://www.uniprot.org/uniprot/Q63TE7) | Drug-resistance related membrane protein | Inner Membrane | Yes | No |
| 203 | [Q63TB3](https://www.uniprot.org/uniprot/Q63TB3) | Uncharacterized protein | Cytoplasmic | No | No |
| 204 | [Q63TB1](https://www.uniprot.org/uniprot/Q63TB1) | Oxidase | Extracellular | No | No |
| 205 | [Q63TB0](https://www.uniprot.org/uniprot/Q63TB0) | Exported protein | Periplasmic | Yes | No |
| 206 | [Q63TA9](https://www.uniprot.org/uniprot/Q63TA9) | SCO family protein | Cytoplasmic | No | No |
| 207 | [Q63TA4](https://www.uniprot.org/uniprot/Q63TA4) | Membrane protein | Extracellular | Yes | No |
| 208 | [Q63TA2](https://www.uniprot.org/uniprot/Q63TA2) | Two component system, response regulator | Cytoplasmic | No | No |
| 209 | [Q63T97](https://www.uniprot.org/uniprot/Q63T97) | Glycoside hydrolase family 44 domain-containing protein | Extracellular | No | No |
| 210 | [Q63T92](https://www.uniprot.org/uniprot/Q63T92) | Maltokinase | Cytoplasmic | No | No |
| 211 | [Q63T89](https://www.uniprot.org/uniprot/Q63T89) | Malto-oligosyltrehalose trehalohydrolase | Extracellular | No | No |
| 212 | [Q63T88](https://www.uniprot.org/uniprot/Q63T88) | 4-alpha-glucanotransferase | Cytoplasmic | No | No |
| 213 | [Q63T87](https://www.uniprot.org/uniprot/Q63T87) | Glycosyl hydrolase | Cytoplasmic | No | No |
| 214 | [Q63T70](https://www.uniprot.org/uniprot/Q63T70) | 2-C-methyl-D-erythritol 4-phosphate cytidylyltransferase (ispD) | Cytoplasmic | No | No |
| 215 | [Q63T50](https://www.uniprot.org/uniprot/Q63T50) | Amino acid racemase | Cytoplasmic | No | No |
| 216 | [Q63T46](https://www.uniprot.org/uniprot/Q63T46) | tRNA-specific adenosine deaminase (tadA) | Cytoplasmic | No | No |
| 217 | [Q63T42](https://www.uniprot.org/uniprot/Q63T42) | GMP synthase [glutamine-hydrolyzing] (guaA) | Cytoplasmic | No | DB04272 DB00142 DB00130 DB00993 |
| 218 | [Q63T40](https://www.uniprot.org/uniprot/Q63T40) | Inosine-5'-monophosphate dehydrogenase (guaB) | Cytoplasmic | No | DB00688 DB01024 DB00811 |
| 219 | [Q63T35](https://www.uniprot.org/uniprot/Q63T35) | Coenzyme Q-binding protein COQ10 START domain-containing protein | Cytoplasmic | No | No |
| 220 | [Q63T2](https://www.uniprot.org/uniprot/Q63T25)5 | Lipid-A-disaccharide synthase (lpxB) | Cytoplasmic | No | No |
| 221 | [Q63T24](https://www.uniprot.org/uniprot/Q63T24) | Acyl-[acyl-carrier-protein]--UDP-N-acetylglucosamine O-acyltransferase (lpxA) | Cytoplasmic | No | No |
| 222 | [Q63T2](https://www.uniprot.org/uniprot/Q63T22)2 | UDP-3-O-acylglucosamine N-acyltransferase (lpxD) | Cytoplasmic | No | No |
| 223 | [Q63T21](https://www.uniprot.org/uniprot/Q63T21) | Outer membrane protein | Cytoplasmic | No | No |
| 224 | [Q63T20](https://www.uniprot.org/uniprot/Q63T20) | Outer membrane protein assembly factor BamA (bamA) | Outer Membrane | No | No |
| 225 | [Q63T19](https://www.uniprot.org/uniprot/Q63T19) | Zinc metalloprotease | Extracellular | Yes | No |
| 226 | [Q63T18](https://www.uniprot.org/uniprot/Q63T18) | 1-deoxy-D-xylulose 5-phosphate reductoisomerase (dxr) | Cytoplasmic | No | No |
| 227 | [Q63T17](https://www.uniprot.org/uniprot/Q63T17) | Phosphatidate cytidylyltransferase | Inner membrane | Yes | No |
| 228 | [Q63T16](https://www.uniprot.org/uniprot/Q63T16) | Isoprenyl transferase (uppS) | Cytoplasmic | No | No |
| 229 | [Q63T14](https://www.uniprot.org/uniprot/Q63T14) | Uridylate kinase (pyrH) | Cytoplasmic | No | No |
| 230 | [Q63T13](https://www.uniprot.org/uniprot/Q63T13) | Elongation factor Ts (tsf) | Cytoplasmic | No | No |
| 231 | [Q63T0](https://www.uniprot.org/uniprot/Q63T06)6 | Cell division protein ZipA | Cytoplasmic | Yes | No |
| 232 | [Q63T05](https://www.uniprot.org/uniprot/Q63T05) | Chromosome partition protein Smc (smc) | Cytoplasmic | No | No |
| 233 | [Q63T02](https://www.uniprot.org/uniprot/Q63T02) | 2,3,4,5-tetrahydropyridine-2,6-dicarboxylate N-succinyltransferase (dapD) | Cytoplasmic | No | No |
| 234 | [Q63T00](https://www.uniprot.org/uniprot/Q63T00) | Succinyl-diaminopimelate desuccinylase (dapE) | Cytoplasmic | No | No |
| 235 | [Q63SZ1](https://www.uniprot.org/uniprot/Q63SZ1) | Membrane protein | Inner membrane | Yes | No |
| 236 | [Q63SX4](https://www.uniprot.org/uniprot/Q63SX4) | Glutamate--tRNA ligase (gltX) | Cytoplasmic | No | DB00130 |
| 237 | [Q63SV](https://www.uniprot.org/uniprot/Q63SV4)4 | Sulfate adenylyltransferase subunit 2 | Cytoplasmic | No | No |
| 238 | [Q63ST2](https://www.uniprot.org/uniprot/Q63ST2) | Aspartokinase (ask) | Cytoplasmic | No | No |
| 239 | [Q63ST1](https://www.uniprot.org/uniprot/Q63ST1) | tRNA(Ile)-lysidine synthase (tilS) | Cytoplasmic | No | No |
| 240 | [Q63ST](https://www.uniprot.org/uniprot/Q63ST0)0 | Acetyl-coenzyme A carboxylase carboxyl transferase subunit alpha (accA) | Cytoplasmic | No | No |
| 241 | [Q63SS8](https://www.uniprot.org/uniprot/Q63SS8) | Cysteine--tRNA ligase (cysS) | Cytoplasmic | No | DB00151 |
| 242 | [Q63SS](https://www.uniprot.org/uniprot/Q63SS4)4 | UDP-2,3-diacylglucosamine hydrolase (lpxH) | Cytoplasmic | No | No |
| 243 | [Q63SS1](https://www.uniprot.org/uniprot/Q63SS1) | Inositol-1-monophosphatase (suhB) | Cytoplasmic | No | DB14507 DB14509 DB01356 DB14508 |
| 244 | [Q63SR2](https://www.uniprot.org/uniprot/Q63SR2) | 4-hydroxy-tetrahydrodipicolinate synthase (dapA) | Cytoplasmic | No | No |
| 245 | [Q63SR0](https://www.uniprot.org/uniprot/Q63SR0) | Tryptophan--tRNA ligase (trpS) | Cytoplasmic | No | DB00150 |
| 246 | [Q63SQ0](https://www.uniprot.org/uniprot/Q63SQ0) | Enolase (eno) | Cytoplasmic | No | No |
| 247 | [Q63SP3](https://www.uniprot.org/uniprot/Q63SP3) | Lipoprotein releasing system transmembrane protein | Inner Membrane | Yes | No |
| 248 | [Q63SP0](https://www.uniprot.org/uniprot/Q63SP0) | Peptide chain release factor 2 (prfB) | Cytoplasmic | No | No |
| 249 | [Q63SN](https://www.uniprot.org/uniprot/Q63SN9)9 | Lysine--tRNA ligase (lysS) | Cytoplasmic | No | DB00123 |
| 250 | [Q63SN1](https://www.uniprot.org/uniprot/Q63SN1) | Cysteine desulfurase IscS (iscS) | Cytoplasmic | No | DB00151 |
| 251 | [Q63SL6](https://www.uniprot.org/uniprot/Q63SL6) | Bifunctional protein FolD (folD) | Cytoplasmic | No | DB00116 DB00157 |
| 252 | [Q63SK9](https://www.uniprot.org/uniprot/Q63SK9) | Respiratory nitrate reductase delta chain | Cytoplasmic | No | No |
| 253 | [Q63SK](https://www.uniprot.org/uniprot/Q63SK2)2 | Glutamine synthetase (glnA) | Cytoplasmic | No | DB04272 DB11638 |
| 254 | [Q63SH3](https://www.uniprot.org/uniprot/Q63SH3) | LuxR family regulatory protein | Cytoplasmic | No | No |
| 255 | [Q63SH0](https://www.uniprot.org/uniprot/Q63SH0) | Membrane protein | Inner Membrane | Yes | No |
| 256 | [Q63SG](https://www.uniprot.org/uniprot/Q63SG3)3 | Ubiquinone biosynthesis accessory factor UbiT (ubiT) | Inner Membrane | No | No |
| 257 | [Q63SG2](https://www.uniprot.org/uniprot/Q63SG2) | Ubiquinone biosynthesis protein UbiV (ubiV) | Cytoplasmic | No | No |
| 258 | [Q63SG0](https://www.uniprot.org/uniprot/Q63SG0) | 2-nitropropane dioxygenase | Cytoplasmic | No | No |
| 259 | [Q63SF9](https://www.uniprot.org/uniprot/Q63SF9) | Membrane protein | Inner membrane | Yes | No |
| 260 | [Q63SF7](https://www.uniprot.org/uniprot/Q63SF7) | Coproporphyrinogen-III oxidase | Cytoplasmic | No | No |
| 261 | [Q63SF](https://www.uniprot.org/uniprot/Q63SF5)5 | Membrane protein | Inner membrane | Yes | No |
| 262 | [Q63SF1](https://www.uniprot.org/uniprot/Q63SF1) | cysteine desulfurase | Cytoplasmic | No | DB11135 |
| 263 | [Q63SE](https://www.uniprot.org/uniprot/Q63SE9)9 | MIP18 family-like domain-containing protein | Cytoplasmic | No | No |
| 264 | [Q63SE](https://www.uniprot.org/uniprot/Q63SE5)5 | Ubiquinol oxidase subunit 2 (cyoA) | Inner membrane | Yes | No |
| 265 | [Q63SA](https://www.uniprot.org/uniprot/Q63SA5)5 | CDP-diacylglycerol--glycerol-3-phosphate 3-phosphatidyltransferase | Inner membrane | Yes | No |
| 266 | [Q63SA2](https://www.uniprot.org/uniprot/Q63SA2) | Elongation factor P (efp) | Cytoplasmic | No | No |
| 267 | [Q3V7S9](https://www.uniprot.org/uniprot/Q3V7S9) | Pyridoxine 5'-phosphate synthase (pdxJ) | Cytoplasmic | No | No |
| 268 | [Q63S89](https://www.uniprot.org/uniprot/Q63S89) | RNA polymerase sigma factor (rpoE) | Cytoplasmic | No | No |
| 269 | [Q63S87](https://www.uniprot.org/uniprot/Q63S87) | 3-oxoacyl-[acyl-carrier-protein] synthase 2 (fabF) | Cytoplasmic | No | DB01034 DB03017 |
| 270 | [Q63S8](https://www.uniprot.org/uniprot/Q63S85)5 | 3-oxoacyl-[acyl-carrier-protein] reductase (fabG) | Cytoplasmic | No | No |
| 271 | [Q63S83](https://www.uniprot.org/uniprot/Q63S83) | Beta-ketoacyl-[acyl-carrier-protein] synthase III (fabH) | Cytoplasmic | No | DB03017 DB03264 DB07650 |
| 272 | [Q63S82](https://www.uniprot.org/uniprot/Q63S82) | Phosphate acyltransferase (plsX) | Cytoplasmic | No | No |
| 273 | [Q63S5](https://www.uniprot.org/uniprot/Q63S51)1 | Thymidylate synthase (thyA) | Cytoplasmic | No | DB00293 DB00322 DB00544 DB00642 DB01101 DB00432 DB01099 |
| 274 | [Q63S46](https://www.uniprot.org/uniprot/Q63S46) | Dihydrofolate reductase | Cytoplasmic | No | DB00440 DB00951 DB00563 DB01157 DB00642 DB01131 DB03904 |
| 275 | [Q63S45](https://www.uniprot.org/uniprot/Q63S45) | Transposase | Cytoplasmic | No | No |
| 276 | [Q63R56](https://www.uniprot.org/uniprot/Q63R56) | Transposase (tnpA) | Cytoplasmic | No | No |
| 277 | [Q63S31](https://www.uniprot.org/uniprot/Q63S31) | Ribosome maturation factor RimM (rimM) | Cytoplasmic | No | No |
| 278 | [Q63S15](https://www.uniprot.org/uniprot/Q63S15) | Cysteine synthase (cysM) | Cytoplasmic | No | No |
| 279 | [P0DMK5](https://www.uniprot.org/uniprot/P0DMK5) | ADP-L-glycero-D-manno-heptose-6-epimerase (hldD) | Cytoplasmic | No | No |
| 280 | [H7C754](https://www.uniprot.org/uniprot/H7C754) | ADP-heptose synthase (waaE) | Cytoplasmic | No | No |
| 281 | [Q63S1](https://www.uniprot.org/uniprot/Q63S10)0 | UDP-glucose 6-dehydrogenase (udg) | Cytoplasmic | No | DB00157 DB09130 |
| 282 | [Q63S0](https://www.uniprot.org/uniprot/Q63S09)9 | Lipopolysaccharide assembly protein B (lapB) | Cytoplasmic | Yes | No |
| 283 | [Q63S0](https://www.uniprot.org/uniprot/Q63S06)6 | 30S ribosomal protein S1 (rpsA) | Cytoplasmic | No | No |
| 284 | [Q63S05](https://www.uniprot.org/uniprot/Q63S05) | Cytidylate kinase (cmk) | Cytoplasmic | No | No |
| 285 | [Q63S01](https://www.uniprot.org/uniprot/Q63S01) | Exported protein | Periplasmic | No | No |
| 286 | [Q63S00](https://www.uniprot.org/uniprot/Q63S00) | DNA gyrase subunit A (gyrA) | Cytoplasmic | No | DB00537 DB11943 DB00487 DB00218 DB00467 DB06771 DB01044 |
| 287 | [Q63RZ](https://www.uniprot.org/uniprot/Q63RZ9)9 | Outer membrane protein a (ompA) | Periplasmic | No | No |
| 288 | [Q63RZ8](https://www.uniprot.org/uniprot/Q63RZ8) | Ubiquinone biosynthesis O-methyltransferase (ubiG) | Cytoplasmic | No | No |
| 289 | [Q63RV9](https://www.uniprot.org/uniprot/Q63RV9) | Guanosine-3',5'-bis(Diphosphate) 3'-pyrophosphohydrolase (spoT) | Cytoplasmic | No | No |
| 290 | [Q63RV7](https://www.uniprot.org/uniprot/Q63RV7) | Guanylate kinase (gmk) | Cytoplasmic | No | DB01972 DB00577 DB00787 |
| 291 | [Q63RS1](https://www.uniprot.org/uniprot/Q63RS1) | Serine--tRNA ligase (serS) | Cytoplasmic | No | No |
| 292 | [Q63RP8](https://www.uniprot.org/uniprot/Q63RP8) | Glutamate-1-semialdehyde 2,1-aminomutase (hemL) | Cytoplasmic | No | No |
| 293 | [Q63RP](https://www.uniprot.org/uniprot/Q63RP5)5 | 6,7-dimethyl-8-ribityllumazine synthase (ribH) | Cytoplasmic | No | No |
| 294 | [Q63RM1](https://www.uniprot.org/uniprot/Q63RM1) | Exported protein | Periplasmic | No | No |
| 295 | [Q63RL9](https://www.uniprot.org/uniprot/Q63RL9) | Branched-chain amino acid transport system, permease component | Inner membrane | Yes | No |
| 296 | [Q63RL7](https://www.uniprot.org/uniprot/Q63RL7) | ABC transport ATP-binding subunit | Cytoplasmic | No | No |
| 297 | [Q63RK7](https://www.uniprot.org/uniprot/Q63RK7) | Lipopolysaccharide heptosyltransferase-1 (waaC1) | Cytoplasmic | No | No |
| 298 | [Q63RK6](https://www.uniprot.org/uniprot/Q63RK6) | Phosphoglucomutase (pgm) | Cytoplasmic | No | DB06773 |
| 299 | [Q63RK5](https://www.uniprot.org/uniprot/Q63RK5) | Lipopolysaccharide biosynthesis protein | Inner membrane | Yes | No |
| 300 | [Q63RK4](https://www.uniprot.org/uniprot/Q63RK4) | Glycosyl transferase | Cytoplasmic | No | No |
| 301 | [H7C761](https://www.uniprot.org/uniprot/H7C761) | dTDP-4-dehydrorhamnose 3,5-epimerase (rmlC) | Cytoplasmic | No | No |
| 302 | [Q9F712](https://www.uniprot.org/uniprot/Q9F712) | Chaperonin GroEL 1 (groEL1) | Cytoplasmic | No | No |
| 303 | [Q63RG](https://www.uniprot.org/uniprot/Q63RG4)4 | Exported Metallo-beta-lactamase-family protein | Extracellular | No | No |
| 304 | [Q63RB](https://www.uniprot.org/uniprot/Q63RB4)4 | Serine hydroxymethyltransferase 1 (glyA1) | Cytoplasmic | No | DB11596 DB02824 DB00116 DB00145 DB11638 DB01055 |
| 305 | [Q63RA8](https://www.uniprot.org/uniprot/Q63RA8) | Tol-Pal system protein TolB (tolB) | Outer Membrane | No | No |
| 306 | [Q63R99](https://www.uniprot.org/uniprot/Q63R99) | Glycosyltransferase | Cytoplasmic | No | No |
| 307 | [Q63R96](https://www.uniprot.org/uniprot/Q63R96) | O-antigen translocase | Inner Membrane | Yes | No |
| 308 | [Q63R93](https://www.uniprot.org/uniprot/Q63R93) | Capsular polysaccharide transport protein | Outer Membrane | No | No |
| 309 | [Q63R92](https://www.uniprot.org/uniprot/Q63R92) | Transmembrane sugar transferase | Inner Membrane | Yes | No |
| 310 | [Q63R56](https://www.uniprot.org/uniprot/Q63R56) | Transposase (tnpA) | Cytoplasmic | No | No |
| 311 | [Q63R4](https://www.uniprot.org/uniprot/Q63R45)5 | Protein GrpE (grpE) | Cytoplasmic | No | No |
| 312 | [Q63R41](https://www.uniprot.org/uniprot/Q63R41) | NAD kinase (nadK) | Cytoplasmic | No | No |
| 313 | [Q63R03](https://www.uniprot.org/uniprot/Q63R03) | Protein translocase subunit SecD (secD) | Inner Membrane | Yes | No |
| 314 | [Q63QY](https://www.uniprot.org/uniprot/Q63QY6)6 | Phosphatase NudJ (nudJ) | Cytoplasmic | No | No |
| 315 | [Q63QY1](https://www.uniprot.org/uniprot/Q63QY1) | Ubiquinone biosynthesis-related protein | Cytoplasmic | No | No |
| 316 | [Q63QX6](https://www.uniprot.org/uniprot/Q63QX6) | Holliday junction branch migration complex subunit RuvA (ruvA) | Cytoplasmic | No | No |
| 317 | [Q63QX5](https://www.uniprot.org/uniprot/Q63QX5) | Holliday junction branch migration complex subunit RuvB (ruvB) | Cytoplasmic | No | DB00173 |
| 318 | [Q63QX0](https://www.uniprot.org/uniprot/Q63QX0) | Tyrosine--tRNA ligase (tyrS) | Cytoplasmic | No | DB00135 |
| 319 | [Q63QW3](https://www.uniprot.org/uniprot/Q63QW3) | Large ribosomal subunit protein uL13 (rplM) | Cytoplasmic | No | No |
| 320 | [Q63QW0](https://www.uniprot.org/uniprot/Q63QW0) | Dihydroorotase (pyrC) | Cytoplasmic | No | No |
| 321 | [Q63QU8](https://www.uniprot.org/uniprot/Q63QU8) | LysR-family transcriptional regulator | Cytoplasmic | No | No |
| 322 | [Q63QU6](https://www.uniprot.org/uniprot/Q63QU6) | Adenylosuccinate lyase (purB) | Cytoplasmic | No | No |
| 323 | [Q63QT6](https://www.uniprot.org/uniprot/Q63QT6) | Leucine--tRNA ligase (leuS) | Cytoplasmic | No | No |
| 324 | [Q63QS1](https://www.uniprot.org/uniprot/Q63QS1) | transketolase (tktA) | Cytoplasmic | No | DB01987 DB09130 |
| 325 | [Q63QR4](https://www.uniprot.org/uniprot/Q63QR4) | Thiamine-monophosphate kinase (thiL) | Cytoplasmic | No | No |
| 326 | [Q63QP0](https://www.uniprot.org/uniprot/Q63QP0) | Biotin carboxylase (accC) | Cytoplasmic | No | DB00121 DB00173 |
| 327 | [Q63QN3](https://www.uniprot.org/uniprot/Q63QN3) | Ribonucleoside-diphosphate reductase subunit beta (nrdB) | Cytoplasmic | Yes | DB09462 |
| 328 | [Q63QM8](https://www.uniprot.org/uniprot/Q63QM8) | Signal recognition particle protein (ffh) | Inner membrane | No | No |
| 329 | [Q63QM](https://www.uniprot.org/uniprot/Q63QM5)5 | Proline--tRNA ligase (proS) | Cytoplasmic | No | No |
| 330 | [Q63QM2](https://www.uniprot.org/uniprot/Q63QM2) | Transposase | Cytoplasmic | No | No |
| 331 | [Q63QM1](https://www.uniprot.org/uniprot/Q63QM1) | Glutamate 5-kinase (proB) | Cytoplasmic | No | No |
| 332 | [Q63QM](https://www.uniprot.org/uniprot/Q63QM0)0 | GTPase Obg (obg) | Cytoplasmic | No | No |
| 333 | [Q63QL7](https://www.uniprot.org/uniprot/Q63QL7) | Octaprenyl-diphosphate synthase (ispB) | Cytoplasmic | No | No |
| 334 | [Q63QK4](https://www.uniprot.org/uniprot/Q63QK4) | UDP-3-O-acyl-N-acetylglucosamine deacetylase (lpxC) | Cytoplasmic | No | No |
| 335 | [Q63QK](https://www.uniprot.org/uniprot/Q63QK2)2 | Cell division protein FtsZ (ftsZ) | Cytoplasmic | No | No |
| 336 | [Q63QK1](https://www.uniprot.org/uniprot/Q63QK1) | Cell division protein FtsA (ftsA) | Cytoplasmic | No | No |
| 337 | [Q63QK0](https://www.uniprot.org/uniprot/Q63QK0) | Cell division protein FtsQ (ftsQ) | Periplasmic | Yes | No |
| 338 | [Q63QJ9](https://www.uniprot.org/uniprot/Q63QJ9) | D-alanine--D-alanine ligase (ddl) | Cytoplasmic | No | DB00260 |
| 339 | [Q63QJ7](https://www.uniprot.org/uniprot/Q63QJ7) | UDP-N-acetylglucosamine--N-acetylmuramyl-(pentapeptide) pyrophosphoryl-undecaprenol N-acetylglucosamine transferase (murG) | Cytoplasmic | No | No |
| 340 | [Q63QJ6](https://www.uniprot.org/uniprot/Q63QJ6) | Probable peptidoglycan glycosyltransferase FtsW (ftsW) | Inner Membrane | Yes | No |
| 341 | [Q63QJ5](https://www.uniprot.org/uniprot/Q63QJ5) | UDP-N-acetylmuramoylalanine--D-glutamate ligase (murD) | Cytoplasmic | No | No |
| 342 | [Q63QJ](https://www.uniprot.org/uniprot/Q63QJ4)4 | Phospho-N-acetylmuramoyl-pentapeptide-transferase (mraY) | Inner Membrane | Yes | No |
| 343 | [Q63QJ3](https://www.uniprot.org/uniprot/Q63QJ3) | UDP-N-acetylmuramoyl-tripeptide--D-alanyl-D-alanine ligase (murF) | Cytoplasmic | No | No |
| 344 | [Q63QJ2](https://www.uniprot.org/uniprot/Q63QJ2) | UDP-N-acetylmuramoyl-L-alanyl-D-glutamate--2,6-diaminopimelate ligase (murE) | Cytoplasmic | No | No |
| 345 | [Q63QJ](https://www.uniprot.org/uniprot/Q63QJ1)1 | Peptidoglycan D,D-transpeptidase FtsI (ftsI) | Periplasmic | Yes | DB00267 DB01416 DB01329 DB01327 DB01331 DB01328 DB01413 DB01415 DB00430 DB00303 |
| 346 | [Q63QI9](https://www.uniprot.org/uniprot/Q63QI9) | Ribosomal RNA small subunit methyltransferase H (rsmH) | Cytoplasmic | No | No |
| 347 | [Q63QI7](https://www.uniprot.org/uniprot/Q63QI7) | 3-demethoxyubiquinol 3-hydroxylase (coq7) | Cytoplasmic | No | No |
| 348 | [Q63QB0](https://www.uniprot.org/uniprot/Q63QB0) | Transcriptional regulator | Cytoplasmic | No | No |
| 349 | [Q63QA](https://www.uniprot.org/uniprot/Q63QA8)8 | Integrase catalytic domain-containing protein | Cytoplasmic | No | No |
| 350 | [Q63QA5](https://www.uniprot.org/uniprot/Q63QA5) | Stringent starvation protein A | Cytoplasmic | No | DB04272 |
| 351 | [Q63Q44](https://www.uniprot.org/uniprot/Q63Q44) | Cytochrome C biogenesis protein | Inner membrane | Yes | No |
| 352 | [Q63Q37](https://www.uniprot.org/uniprot/Q63Q37) | DNA-directed RNA polymerase subunit alpha (rpoA) | Cytoplasmic | No | DB00615 DB11753 |
| 353 | [Q63Q36](https://www.uniprot.org/uniprot/Q63Q36) | Small ribosomal subunit protein uS4 (rpsD) | Cytoplasmic | No | DB00453 DB00595 DB01017 DB12329 DB00256 |
| 354 | [Q63Q31](https://www.uniprot.org/uniprot/Q63Q31) | Protein translocase subunit SecY (secY) | Inner membrane | Yes | No |
| 355 | [Q63Q28](https://www.uniprot.org/uniprot/Q63Q28) | Small ribosomal subunit protein uS5 (rpsE) | Cytoplasmic | No | No |
| 356 | [Q63Q26](https://www.uniprot.org/uniprot/Q63Q26) | Large ribosomal subunit protein uL6 (rplF) | Cytoplasmic | No | No |
| 357 | [Q63Q23](https://www.uniprot.org/uniprot/Q63Q23) | Large ribosomal subunit protein uL5 (rplE) | Cytoplasmic | No | No |
| 358 | [Q63Q1](https://www.uniprot.org/uniprot/Q63Q17)7 | Small ribosomal subunit protein uS3 (rpsC) | Cytoplasmic | No | DB00759 DB09093 DB12455 |
| 359 | [Q63Q14](https://www.uniprot.org/uniprot/Q63Q14) | Large ribosomal subunit protein uL2 (rplB) | Cytoplasmic | No | DB04865 |
| 360 | [Q63Q12](https://www.uniprot.org/uniprot/Q63Q12) | Large ribosomal subunit protein uL4 (rplD) | Cytoplasmic | No | DB13179 DB04272 |
| 361 | [Q63Q11](https://www.uniprot.org/uniprot/Q63Q11) | Large ribosomal subunit protein uL3 (rplC) | Cytoplasmic | No | DB01256 DB06145 |
| 362 | [Q63PZ6](https://www.uniprot.org/uniprot/Q63PZ6) | Elongation factor Tu (tuf) | Cytoplasmic | No | DB04315 DB01593 DB14487 DB14533 |
| 363 | [Q63Q08](https://www.uniprot.org/uniprot/Q63Q08) | Elongation factor G 2 (fusA2) | Cytoplasmic | No | DB04315 DB02703 |
| 364 | [Q63Q07](https://www.uniprot.org/uniprot/Q63Q07) | Small ribosomal subunit protein uS7 (rpsG) | Cytoplasmic | No | DB00759 DB13092 DB09093 DB12455 |
| 365 | [Q63Q0](https://www.uniprot.org/uniprot/Q63Q04)4 | DNA-directed RNA polymerase subunit beta' (rpoC) | Cytoplasmic | No | DB00615 DB11753 DB01201 |
| 366 | [Q63Q03](https://www.uniprot.org/uniprot/Q63Q03) | DNA-directed RNA polymerase subunit beta (rpoB) | Cytoplasmic | No | DB01045 DB01220 DB04934 DB00615 DB11753 |
| 367 | [Q63Q01](https://www.uniprot.org/uniprot/Q63Q01) | Large ribosomal subunit protein uL10 (rplJ) | Cytoplasmic | No | DB00778 DB01211 DB01369 |
| 368 | [Q63PZ8](https://www.uniprot.org/uniprot/Q63PZ8) | Transcription termination/antitermination protein NusG (nusG) | Cytoplasmic | No | No |
| 369 | [Q63PZ6](https://www.uniprot.org/uniprot/Q63PZ6) | Elongation factor Tu (tuf) | Cytoplasmic | No | DB04315 DB01593 DB14487 DB14533 |
| 370 | [Q63PI6](https://www.uniprot.org/uniprot/Q63PI6) | Primosomal protein N' (priA) | Cytoplasmic | No | No |
| 371 | [Q63PI](https://www.uniprot.org/uniprot/Q63PI0)0 | ATP synthase subunit beta 1 (atpD1) | Cytoplasmic | No | DB01119 DB04216 DB08949 |
| 372 | [Q63PH9](https://www.uniprot.org/uniprot/Q63PH9) | ATP synthase gamma chain (atpG) | Cytoplasmic | No | DB01119 DB04216 |
| 373 | [Q63PH8](https://www.uniprot.org/uniprot/Q63PH8) | ATP synthase subunit alpha 1 (atpA1) | Inner Membrane | No | DB11638 DB01119 DB04216 |
| 374 | [Q63PH](https://www.uniprot.org/uniprot/Q63PH6)6 | ATP synthase subunit b (atpF) | Inner Membrane | No | No |
| 375 | [Q63PH4](https://www.uniprot.org/uniprot/Q63PH4) | ATP synthase subunit a (atpB) | Inner Membrane | Yes | No |
| 376 | [Q63PH3](https://www.uniprot.org/uniprot/Q63PH3) | ATP synthase protein I AtpI | Inner Membrane | Yes | No |
| 377 | [Q63PH1](https://www.uniprot.org/uniprot/Q63PH1) | Chromosome partitioning protein ParB | Cytoplasmic | No | No |
| 378 | [Q63P53](https://www.uniprot.org/uniprot/Q63P53) | Outer membrane usher protein | Outer Membrane | No | No |
| 379 | [Q63P48](https://www.uniprot.org/uniprot/Q63P48) | Type VI secretion system contractile sheath large subunit | Cytoplasmic | No | No |
| 380 | [Q63P41](https://www.uniprot.org/uniprot/Q63P41) | Gp5/Type VI secretion system Vgr protein OB-fold domain-containing protein | Extracellular | No | No |
| 381 | [Q63P36](https://www.uniprot.org/uniprot/Q63P36) | Type VI secretion system baseplate subunit TssF | Cytoplasmic | No | No |
| 382 | [Q63P32](https://www.uniprot.org/uniprot/Q63P32) | Chaperone-related protein | Cytoplasmic | No | No |
| 383 | [Q63P00](https://www.uniprot.org/uniprot/Q63P00) | Subfamily M20D non-peptidase homologue | Cytoplasmic | No | No |
| 384 | [Q63NY](https://www.uniprot.org/uniprot/Q63NY5)5 | Asparagine synthetase [glutamine-hydrolyzing] (asnO) | Cytoplasmic | No | DB00128 DB00171 DB00174 DB00142 DB00130 |
| 385 | [Q63NP8](https://www.uniprot.org/uniprot/Q63NP8) | Porin membrane protein | Outer Membrane | No | No |
| 386 | [Q63N9](https://www.uniprot.org/uniprot/Q63N94)4 | Bacteriophage/transposase fusion protein | Cytoplasmic | No | No |
| 387 | [Q63MY1](https://www.uniprot.org/uniprot/Q63MY1) | EvpB family type VI secretion protein | Cytoplasmic | No | No |
| 388 | [Q63MX4](https://www.uniprot.org/uniprot/Q63MX4) | Gp5/Type VI secretion system Vgr protein OB-fold domain-containing protein | Outer Membrane | No | No |
| 389 | [Q63MW1](https://www.uniprot.org/uniprot/Q63MW1) | UDP-glucoronosyl and UDP-glucosyl transferase | Cytoplasmic | No | No |
| 390 | [Q63MV3](https://www.uniprot.org/uniprot/Q63MV3) | Glutathione-independent formaldehyde dehydrogenase (fdhA) | Cytoplasmic | No | No |
| 391 | [Q63MU7](https://www.uniprot.org/uniprot/Q63MU7) | Membrane protein | Inner Membrane | Yes | No |
| 392 | [Q63MP1](https://www.uniprot.org/uniprot/Q63MP1) | Thiamine pyrophosphate enzyme family protein | Cytoplasmic | No | No |
| 393 | [Q63MH4](https://www.uniprot.org/uniprot/Q63MH4) | SfnB family sulfur acquisition oxidoreductase | Cytoplasmic | No | No |
| 394 | [Q63MH3](https://www.uniprot.org/uniprot/Q63MH3) | SfnB family sulfur acquisition oxidoreductase | Cytoplasmic | No | No |
| 395 | [Q63MC5](https://www.uniprot.org/uniprot/Q63MC5) | Glycosyl transferase | Inner Membrane | Yes | No |
| 396 | [Q63MB4](https://www.uniprot.org/uniprot/Q63MB4) | Lipase | Periplasmic | No | No |
| 397 | [Q63MB0](https://www.uniprot.org/uniprot/Q63MB0) | Amino acid permease | Inner Membrane | Yes | No |
| 398 | [Q63MA8](https://www.uniprot.org/uniprot/Q63MA8) | Lipoprotein | Periplasmic | No | No |
| 399 | [Q63MA6](https://www.uniprot.org/uniprot/Q63MA6) | Membrane protein | Inner Membrane | Yes | No |
| 400 | [Q63M80](https://www.uniprot.org/uniprot/Q63M80) | Sugar transport protein | Inner Membrane | Yes | No |
| 401 | [Q63M66](https://www.uniprot.org/uniprot/Q63M66) | Surface-exposed protein | Extracellular | No | No |
| 402 | [Q63M47](https://www.uniprot.org/uniprot/Q63M47) | histidine kinase | Inner Membrane | No | No |
| 403 | [Q63M37](https://www.uniprot.org/uniprot/Q63M37) | Glutathione synthetase | Inner Membrane | No | No |
| 404 | [Q63LZ4](https://www.uniprot.org/uniprot/Q63LZ4) | Phenylacetaldehyde dehydrogenase (feaB) | Cytoplasmic | No | No |
| 405 | [Q63LZ3](https://www.uniprot.org/uniprot/Q63LZ3) | Aminotransferase class-III | Cytoplasmic | No | No |
| 406 | [Q63LH8](https://www.uniprot.org/uniprot/Q63LH8) | Sugar transport protein | Inner Membrane | Yes | No |
| 407 | [Q63LH3](https://www.uniprot.org/uniprot/Q63LH3) | Cobalt-zinc-cadmium resistance protein (czcB) | Inner Membrane | Yes | No |
| 408 | [Q63LC7](https://www.uniprot.org/uniprot/Q63LC7) | Bacteriophage protein gp17 | Extracellular | No | No |
| 409 | [Q63LA8](https://www.uniprot.org/uniprot/Q63LA8) | Cation transport ATPase protein | Inner Membrane | Yes | DB11638 |
| 410 | [Q63L78](https://www.uniprot.org/uniprot/Q63L78) | Uncharacterized protein | Cytoplasmic | No | No |
| 411 | [Q63L65](https://www.uniprot.org/uniprot/Q63L65) | Formate hydrogenlyase/hydrogenase/NADH dehydrogenase subunit | Cytoplasmic | No | No |
| 412 | [Q63L44](https://www.uniprot.org/uniprot/Q63L44) | Xaa-Pro dipeptidyl-peptidase C-terminal domain-containing protein | Periplasmic | No | No |
| 413 | [Q63L42](https://www.uniprot.org/uniprot/Q63L42) | Transport/efflux protein | Inner Membrane | Yes | No |
| 414 | [Q63KZ4](https://www.uniprot.org/uniprot/Q63KZ4) | Peptidoglycan D,D-transpeptidase FtsI (ftsI) | Inner Membrane | Yes | No |
| 415 | [Q63KY9](https://www.uniprot.org/uniprot/Q63KY9) | Exported protein | Cytoplasmic | Yes | No |
| 416 | [Q63KX7](https://www.uniprot.org/uniprot/Q63KX7) | Sugar transporter ATP-binding protein | Inner Membrane | No | No |
| 417 | [Q63KU2](https://www.uniprot.org/uniprot/Q63KU2) | Acyl-CoA dehydrogenase | Cytoplasmic | No | DB03059 DB03147 DB00157 |
| 418 | [Q63KN0](https://www.uniprot.org/uniprot/Q63KN0) | Phosphatase protein | Periplasmic | No | No |
| 419 | [Q63KM0](https://www.uniprot.org/uniprot/Q63KM0) | cysteine desulfurase | Cytoplasmic | No | DB00160 DB00151 |
| 420 | [Q63KL1](https://www.uniprot.org/uniprot/Q63KL1) | Efflux/sugar transport/multidrug resistance protein | Inner Membrane | Yes | No |
| 421 | [Q63KL0](https://www.uniprot.org/uniprot/Q63KL0) | Rhamnosyltransferase protein | Cytoplasmic | No | No |
| 422 | [Q63KK4](https://www.uniprot.org/uniprot/Q63KK4) | histidine kinase | Cytoplasmic | No | No |
| 423 | [Q63KI0](https://www.uniprot.org/uniprot/Q63KI0) | Transmembrane phospholipase protein | Inner Membrane | Yes | No |
| 424 | [Q63KH2](https://www.uniprot.org/uniprot/Q63KH2) | Membrane protein | Extracellular | Yes | No |
| 425 | [Q63KH0](https://www.uniprot.org/uniprot/Q63KH0) | Type 3 secretion system secretin (sctC) | Outer Membrane | Yes | No |
| 426 | [Q63KG9](https://www.uniprot.org/uniprot/Q63KG9) | AraC family regulator of pathogenicity genes | Cytoplasmic | No | No |
| 427 | [Q63KB2](https://www.uniprot.org/uniprot/Q63KB2) | DUF3326 domain-containing protein | Cytoplasmic | No | No |
| 428 | [Q63K78](https://www.uniprot.org/uniprot/Q63K78) | Copper-containing nitrite reductase | Periplasmic | Yes | No |
| 429 | [Q63K63](https://www.uniprot.org/uniprot/Q63K63) | Clp-type ATPase chaperone protein | Outer Membrane | No | No |
| 430 | [Q63K54](https://www.uniprot.org/uniprot/Q63K54) | Membrane protein | Outer Membrane | Yes | No |
| 431 | [Q63K53](https://www.uniprot.org/uniprot/Q63K53) | Membrane protein | Periplasmic | No | No |
| 432 | [Q63JV4](https://www.uniprot.org/uniprot/Q63JV4) | AraC-family transcriptional regulator | Cytoplasmic | No | No |
| 433 | [Q63JU4](https://www.uniprot.org/uniprot/Q63JU4) | Type III secretion protein | Inner Membrane | Yes | No |
| 434 | [Q63JT2](https://www.uniprot.org/uniprot/Q63JT2) | Probable non-ribosomal peptide synthetase | Cytoplasmic | No | No |
| 435 | [Q63JT1](https://www.uniprot.org/uniprot/Q63JT1) | Probable non-ribosomal peptide synthetase | Cytoplasmic | No | No |
| 436 | [Q63JT0](https://www.uniprot.org/uniprot/Q63JT0) | Probable non-ribosomal peptide synthetase | Cytoplasmic | No | No |
| 437 | [Q63JS0](https://www.uniprot.org/uniprot/Q63JS0) | histidine kinase | Inner Membrane | No | No |
| 438 | [Q63JQ9](https://www.uniprot.org/uniprot/Q63JQ9) | Beta-glucosidase (bglB) | Periplasmic | No | No |
| 439 | [Q63JN](https://www.uniprot.org/uniprot/Q63JN2)2 | Lipopolysaccharide biosynthesis related membrane protein | Inner Membrane | Yes | No |
| 440 | [Q63JK1](https://www.uniprot.org/uniprot/Q63JK1) | Citrate synthase (gltA) | Cytoplasmic | No | DB04272 DB01992 |
| 441 | [Q63JJ8](https://www.uniprot.org/uniprot/Q63JJ8) | Succinate dehydrogenase flavoprotein subunit (sdhA) | Inner Membrane | No | DB04657 DB04795 DB09270 DB00139 |
| 442 | [Q63JJ0](https://www.uniprot.org/uniprot/Q63JJ0) | Aconitate hydratase (citB) | Cytoplasmic | No | DB06757 |
| 443 | [Q63J9](https://www.uniprot.org/uniprot/Q63J91)1 | Membrane protein | Inner Membrane | Yes | No |
| 444 | [Q63J88](https://www.uniprot.org/uniprot/Q63J88) | Exopolysaccharide biosynthesis related tyrosine-protein kinase | Inner Membrane | Yes | No |
| 445 | [Q63J31](https://www.uniprot.org/uniprot/Q63J31) | Aromatic hydrocarbons catabolism-related reductase | Cytoplasmic | No | No |
| 446 | [Q63J28](https://www.uniprot.org/uniprot/Q63J28) | Aromatic oxygenase | Cytoplasmic | No | No |
| 447 | [Q63IZ1](https://www.uniprot.org/uniprot/Q63IZ1) | Sulfotransferase | Cytoplasmic | No | No |
| 448 | [Q63IY9](https://www.uniprot.org/uniprot/Q63IY9) | Methyl-accepting chemotaxis protein | Inner Membrane | Yes | No |
| 449 | [Q63IU5](https://www.uniprot.org/uniprot/Q63IU5) | Two component system fusin protein | Cytoplasmic | Yes | No |
| 450 | [Q63IT2](https://www.uniprot.org/uniprot/Q63IT2) | Nucleoside phosphorylase domain-containing protein | Periplasmic | No | No |
| 451 | [Q63IP](https://www.uniprot.org/uniprot/Q63IP5)5 | Zinc-containing alcohol dehydrogenase | Cytoplasmic | No | No |
| 452 | [Q63IH8](https://www.uniprot.org/uniprot/Q63IH8) | Type VI secretion system baseplate subunit TssG | Periplasmic | No | No |
| 453 | [Q63IH4](https://www.uniprot.org/uniprot/Q63IH4) | EvpB family type VI secretion protein | Cytoplasmic | No | No |
| 454 | [Q63I10](https://www.uniprot.org/uniprot/Q63I10) | Fusion protein, ATP-binding transmembrane ABC transporter and regulatory protein | Inner Membrane | Yes | DB05154 |

**Supplementary table 4:** List of extracellular proteins of *Burkholderia pseudomallei* K96243 with VaxiJen v2.0 provided antigenicity value

| **Sl No.** | **Protein** | **UniProt ID** | **VaxiJen Value** | **Antigenicity** | **Pathway** |
| --- | --- | --- | --- | --- | --- |
| 1 | Single-stranded DNA-binding protein (ssb) | [Q63XJ3](https://www.uniprot.org/uniprot/Q63XJ3) | 1.4366 | Yes | bps03030 DNA replication bps03430 Mismatch repair bps03440 Homologous recombination |
| 2 | N-acetylmuramoyl-L-alanine amidase | [Q63WN2](https://www.uniprot.org/uniprot/Q63WN2) | 0.5931 | Yes | bps01503 Cationic antimicrobial peptide (CAMP) resistance |
| 3 | Probable lipid II flippase MurJ (murJ) | [Q63WL9](https://www.uniprot.org/uniprot/Q63WL9) | 0.432 | Yes | No |
| 4 | Superoxide dismutase (sodB) | [Q63WL1](https://www.uniprot.org/uniprot/Q63WL1) | 0.4647 | Yes | No |
| 5 | Ribonuclease H (rnhA) | [Q63V99](https://www.uniprot.org/uniprot/Q63V99) | 0.4693 | Yes | bps03030 DNA replication |
| 6 | Subfamily M23B unassigned peptidase | [Q63UU2](https://www.uniprot.org/uniprot/Q63UU2) | 0.5649 | Yes | No |
| 7 | Nucleoside diphosphate kinase (ndk) | [Q63UT6](https://www.uniprot.org/uniprot/Q63UT6) | 0.5435 | Yes | bps00230 Purine metabolism bps00240 Pyrimidine metabolism bps01100 Metabolic pathways bps01110 Biosynthesis of secondary metabolites bps01232 Nucleotide metabolism bps01240 Biosynthesis of cofactors |
| 8 | Exported fimbria-related protein | [Q63UH3](https://www.uniprot.org/uniprot/Q63UH3) | 0.6819 | Yes | No |
| 9 | Outer membrane protein | [Q63UH1](https://www.uniprot.org/uniprot/Q63UH1) | 1.2837 | Yes | No |
| 10 | Hemolysin-related protein | [Q63UE4](https://www.uniprot.org/uniprot/Q63UE4) | 0.7806 | Yes | No |
| 11 | DUF3443 domain-containing protein | [Q63UD3](https://www.uniprot.org/uniprot/Q63UD3) | 0.6375 | Yes | No |
| 12 | Family M20 unassigned peptidase | [Q63UC4](https://www.uniprot.org/uniprot/Q63UC4) | 0.463 | Yes | bps00240 Pyrimidine metabolism bps00410 beta-Alanine metabolism bps00770 Pantothenate and CoA biosynthesis bps01100 Metabolic pathways |
| 13 | Membrane protein | [Q63UB0](https://www.uniprot.org/uniprot/Q63UB0) | 0.9678 | Yes | No |
| 14 | Exported protein | [Q63UA7](https://www.uniprot.org/uniprot/Q63UA7) | 0.4796 | Yes | No |
| 15 | Exported porin | [Q63U88](https://www.uniprot.org/uniprot/Q63U88) | 0.6857 | Yes | No |
| 16 | Oxidase | [Q63TB1](https://www.uniprot.org/uniprot/Q63TB1) | 0.7333 | Yes | bps00052 Galactose metabolism bps01100 Metabolic pathways |
| 17 | Membrane protein | [Q63TA4](https://www.uniprot.org/uniprot/Q63TA4) | 0.8443 | Yes | No |
| 18 | Glycoside hydrolase family 44 domain-containing protein | [Q63T97](https://www.uniprot.org/uniprot/Q63T97) | 0.7379 | Yes | No |
| 19 | Malto-oligosyltrehalose trehalohydrolase | [Q63T89](https://www.uniprot.org/uniprot/Q63T89) | 0.5017 | Yes | bps00500 Starch and sucrose metabolism bps01100 Metabolic pathways bps01110 Biosynthesis of secondary metabolites |
| 20 | Zinc metalloprotease | [Q63T19](https://www.uniprot.org/uniprot/Q63T19) | 0.6328 | Yes | No |
| 21 | Exported Metallo-beta-lactamase-family protein | [Q63RG](https://www.uniprot.org/uniprot/Q63RG4)4 | 0.4944 | Yes | No |
| 22 | Gp5/Type VI secretion system Vgr protein OB-fold domain-containing protein | [Q63P41](https://www.uniprot.org/uniprot/Q63P41) | 0.8115 | Yes | bps03070 Bacterial secretion system |
| 23 | Surface-exposed protein | [Q63M66](https://www.uniprot.org/uniprot/Q63M66) | 0.9578 | Yes | No |
| 24 | Bacteriophage protein gp17 | [Q63LC7](https://www.uniprot.org/uniprot/Q63LC7) | 0.5418 | Yes | No |
| 25 | Membrane protein | [Q63KH2](https://www.uniprot.org/uniprot/Q63KH2) | 0.6875 | Yes | No |

**Supplementary table 5:** Protein list with Drug Bank IDs.

| **SL No.** | **Protein name** | **UniProt ID** | **Drug Bank** |
| --- | --- | --- | --- |
| 1 | DNA gyrase subunit B | [Q63YW7](https://www.uniprot.org/uniprotkb/Q63YW7/entry) | DB00817 DB04395 DB03966 DB05488 DB01044 DB01051 |
| 2 | ferredoxin-NADP reductase | [Q63YE7](https://www.uniprot.org/uniprot/Q63YE7) | DB03147 |
| 3 | glucosamine--fructose-6-phosphate aminotransferase | [Q63Y76](https://www.uniprot.org/uniprot/Q63Y76) | DB00130 |
| 4 | cell division protein | [Q63XM5](https://www.uniprot.org/uniprot/Q63XM5) | DB04272 |
| 5 | ribose-phosphate pyrophosphokinase | [Q63XL8](https://www.uniprot.org/uniprot/Q63XL8) | DB11638 |
| 6 | aspartyl-tRNA synthetase | [Q63X93](https://www.uniprot.org/uniprot/Q63X93) | DB00128 |
| 7 | UDP-N-acetylenolpyruvoylglucosamine reductase | [Q63WM3](https://www.uniprot.org/uniprot/Q63WM3) | DB03147 DB07296 |
| 8 | isocitrate dehydrogenase | [Q63WJ4](https://www.uniprot.org/uniprot/Q63WJ4) | DB02159 DB04272 DB04349 DB04530 DB00157 DB09130 DB06757 DB09092 |
| 9 | bifunctional phosphopantothenoylcysteine decarboxylasephosphopantothenate synthase | [Q63WI5](https://www.uniprot.org/uniprot/Q63WI5) | DB03738 DB03247 |
| 10 | isoleucyl-tRNA synthetase | [Q63WI3](https://www.uniprot.org/uniprot/Q63WI3) | DB00410 DB11638 |
| 11 | DNA topoisomerase IV subunit B | [Q63W24](https://www.uniprot.org/uniprot/Q63W24) | DB00537 DB01044 |
| 12 | ABC transporter ATP-binding protein | [Q63VX](https://www.uniprot.org/uniprot/Q63VX7)7 | DB05154 |
| 13 | nicotinic acid mononucleotide adenylyltransferase | [Q63VT2](https://www.uniprot.org/uniprot/Q63VT2) | DB04272 |
| 14 | dihydropteroate synthase | [Q63V84](https://www.uniprot.org/uniprot/Q63V84) | DB00576 DB01298 DB00634 DB00259 DB01015 DB01581 DB01582 DB06729 DB00250 DB00664 DB01299 DB01145 DB00359 DB08798 DB06147 |
| 15 | ATP-dependent protease ATP-binding subunit ClpX | [Q63V40](https://www.uniprot.org/uniprot/Q63V40) | DB09275 |
| 16 | nucleoside diphosphate kinase | [Q63UT6](https://www.uniprot.org/uniprot/Q63UT6) | DB04315 DB00709 DB09299 DB01262 DB00787 |
| 17 | ABC transporter ATP-binding protein | [Q63UF4](https://www.uniprot.org/uniprot/Q63UF4) | DB05154 |
| 18 | succinate-semialdehyde dehydrogenase | [Q63UF2](https://www.uniprot.org/uniprot/Q63UF2) | DB00534 DB00157 DB00139 DB00313 DB09072 |
| 19 | argininosuccinate synthase | [Q63U95](https://www.uniprot.org/uniprot/Q63U95) | DB00125 DB00128 DB00155 DB00171 |
| 20 | ABC transporter ATP-binding protein | [Q63TZ4](https://www.uniprot.org/uniprot/Q63TZ4) | DB05154 |
| 21 | 2-oxoglutarate dehydrogenase E1 | [Q63TQ7](https://www.uniprot.org/uniprot/Q63TQ7) | DB00157 DB00313 DB09092 |
| 22 | phenylalanyl-tRNA synthetase subunit beta | [Q63TM7](https://www.uniprot.org/uniprot/Q63TM7) | DB00120 |
| 23 | phenylalanyl-tRNA synthetase subunit alpha | [Q63TM6](https://www.uniprot.org/uniprot/Q63TM6) | DB00120 |
| 24 | threonyl-tRNA synthetase | [Q63TM2](https://www.uniprot.org/uniprot/Q63TM2) | DB00156 DB11638 |
| 25 | valyl-tRNA synthetase | [Q63TI8](https://www.uniprot.org/uniprot/Q63TI8) | DB00161 |
| 26 | ABC transporter ATP-binding protein | [Q63TH](https://www.uniprot.org/uniprot/Q63TH5)5 | DB05154 |
| 27 | glutaminyl-tRNA synthetase | [Q63TF6](https://www.uniprot.org/uniprot/Q63TF6) | DB00130 |
| 28 | GMP synthase | [Q63T42](https://www.uniprot.org/uniprot/Q63T42) | DB04272 DB00142 DB00130 DB00993 |
| 29 | inosine 5'-monophosphate dehydrogenase | [Q63T40](https://www.uniprot.org/uniprot/Q63T40) | DB00688 DB01024 DB00811 |
| 30 | glutamyl-tRNA synthetase | [Q63SX4](https://www.uniprot.org/uniprot/Q63SX4) | DB00130 |
| 31 | cysteinyl-tRNA synthetase | [Q63SS8](https://www.uniprot.org/uniprot/Q63SS8) | DB00151 |
| 32 | inositol monophosphatase | [Q63SS1](https://www.uniprot.org/uniprot/Q63SS1) | DB14507 DB14509 DB01356 DB14508 |
| 33 | tryptophanyl-tRNA synthetase | [Q63SR0](https://www.uniprot.org/uniprot/Q63SR0) | DB00150 |
| 34 | lysyl-tRNA synthetase | [Q63SN](https://www.uniprot.org/uniprot/Q63SN9)9 | DB00123 |
| 35 | cysteine desulfurase | [Q63SN1](https://www.uniprot.org/uniprot/Q63SN1) | DB00151 |
| 36 | bifunctional 5,10-methylene-tetrahydrofolate dehydrogenase 5,10-methylene-tetrahydrofolate cyclohydrolase | [Q63SL6](https://www.uniprot.org/uniprot/Q63SL6) | DB00116 DB00157 |
| 37 | glutamine synthetase | [Q63SK](https://www.uniprot.org/uniprot/Q63SK2)2 | DB04272 DB11638 |
| 38 | selenocysteine lyase | [Q63SF1](https://www.uniprot.org/uniprot/Q63SF1) | DB11135 |
| 39 | 3-oxoacyl-ACP synthase | [Q63S87](https://www.uniprot.org/uniprot/Q63S87) | DB01034 DB03017 |
| 40 | 3-oxoacyl-ACP synthase | [Q63S83](https://www.uniprot.org/uniprot/Q63S83) | DB03017 DB03264 DB07650 |
| 41 | thymidylate synthase | [Q63S5](https://www.uniprot.org/uniprot/Q63S51)1 | DB00293 DB00322 DB00544 DB00642 DB01101 DB00432 DB01099 |
| 42 | dihydrofolate reductase | [Q63S46](https://www.uniprot.org/uniprot/Q63S46) | DB00440 DB00951 DB00563 DB01157 DB00642 DB01131 DB03904 |
| 43 | UDP-glucose dehydrogenase | [Q63S1](https://www.uniprot.org/uniprot/Q63S10)0 | DB00157 DB09130 |
| 44 | DNA gyrase subunit A | [Q63S00](https://www.uniprot.org/uniprot/Q63S00) | DB00537 DB11943 DB00487 DB00218 DB00467 DB06771 DB01044 |
| 45 | guanylate kinase | [Q63RV7](https://www.uniprot.org/uniprot/Q63RV7) | DB01972 DB00577 DB00787 |
| 46 | phosphoglucomutase | [Q63RK6](https://www.uniprot.org/uniprot/Q63RK6) | DB06773 |
| 47 | serine hydroxymethyltransferase | [Q63RB](https://www.uniprot.org/uniprot/Q63RB4)4 | DB11596 DB02824 DB00116 DB00145 DB11638 DB01055 |
| 48 | Holliday junction DNA helicase RuvB | [Q63QX5](https://www.uniprot.org/uniprot/Q63QX5) | DB00173 |
| 49 | tyrosyl-tRNA synthetase | [Q63QX0](https://www.uniprot.org/uniprot/Q63QX0) | DB00135 |
| 50 | transketolase | [Q63QS1](https://www.uniprot.org/uniprot/Q63QS1) | DB01987 DB09130 |
| 51 | acetyl-CoA carboxylase biotin carboxylase subunit | [Q63QP0](https://www.uniprot.org/uniprot/Q63QP0) | DB00121 DB00173 |
| 52 | ribonucleotide-diphosphate reductase subunit beta | [Q63QN3](https://www.uniprot.org/uniprot/Q63QN3) | DB09462 |
| 53 | D-alanine--D-alanine ligase | [Q63QJ9](https://www.uniprot.org/uniprot/Q63QJ9) | DB00260 |
| 54 | peptidoglycan synthetase FtsI | [Q63QJ](https://www.uniprot.org/uniprot/Q63QJ1)1 | DB00267 DB01416 DB01329 DB01327 DB01331 DB01328 DB01413 DB01415 DB00430 DB00303 |
| 55 | stringent starvation protein A | [Q63QA5](https://www.uniprot.org/uniprot/Q63QA5) | DB04272 |
| 56 | DNA-directed RNA polymerase subunit alpha | [Q63Q37](https://www.uniprot.org/uniprot/Q63Q37) | DB00615 DB11753 |
| 57 | 30S ribosomal protein S4 | [Q63Q36](https://www.uniprot.org/uniprot/Q63Q36) | DB00453 DB00595 DB01017 DB12329 DB00256 |
| 58 | 30S ribosomal protein S3 | [Q63Q1](https://www.uniprot.org/uniprot/Q63Q17)7 | DB00759 DB09093 DB12455 |
| 59 | 50S ribosomal protein L2 | [Q63Q14](https://www.uniprot.org/uniprot/Q63Q14) | DB04865 |
| 60 | 50S ribosomal protein L4 | [Q63Q12](https://www.uniprot.org/uniprot/Q63Q12) | DB13179 DB04272 |
| 61 | 50S ribosomal protein L3 | [Q63Q11](https://www.uniprot.org/uniprot/Q63Q11) | DB01256 DB06145 |
| 62 | elongation factor Tu | [Q63PZ6](https://www.uniprot.org/uniprot/Q63PZ6) | DB04315 DB01593 DB14487 DB14533 |
| 63 | elongation factor G | [Q63Q08](https://www.uniprot.org/uniprot/Q63Q08) | DB04315 DB02703 |
| 64 | 30S ribosomal protein S7 | [Q63Q07](https://www.uniprot.org/uniprot/Q63Q07) | DB00759 DB13092 DB09093 DB12455 |
| 65 | DNA-directed RNA polymerase subunit beta' | [Q63Q0](https://www.uniprot.org/uniprot/Q63Q04)4 | DB00615 DB11753 DB01201 |
| 66 | DNA-directed RNA polymerase subunit beta | [Q63Q03](https://www.uniprot.org/uniprot/Q63Q03) | DB01045 DB01220 DB04934 DB00615 DB11753 |
| 67 | 50S ribosomal protein L10 | [Q63Q01](https://www.uniprot.org/uniprot/Q63Q01) | DB00778 DB01211 DB01369 |
| 68 | elongation factor Tu | [Q63PZ6](https://www.uniprot.org/uniprot/Q63PZ6) | DB04315 DB01593 DB14487 DB14533 |
| 69 | ATP synthase F0F1 subunit beta | [Q63PI](https://www.uniprot.org/uniprot/Q63PI0)0 | DB01119 DB04216 DB08949 |
| 70 | ATP synthase F0F1 subunit gamma | [Q63PH9](https://www.uniprot.org/uniprot/Q63PH9) | DB01119 DB04216 |
| 71 | ATP synthase F0F1 subunit alpha | [Q63PH8](https://www.uniprot.org/uniprot/Q63PH8) | DB11638 DB01119 DB04216 |
| 72 | asparagine synthase | [Q63NY](https://www.uniprot.org/uniprot/Q63NY5)5 | DB00128 DB00171 DB00174 DB00142 DB00130 |
| 73 | cation transport ATPase | [Q63LA8](https://www.uniprot.org/uniprot/Q63LA8) | DB11638 |
| 74 | acyl-CoA dehydrogenase | [Q63KU2](https://www.uniprot.org/uniprot/Q63KU2) | DB03059 DB03147 DB00157 |
| 75 | cysteine desulfurase | [Q63KM0](https://www.uniprot.org/uniprot/Q63KM0) | DB00160 DB00151 |
| 76 | type II citrate synthase | [Q63JK1](https://www.uniprot.org/uniprot/Q63JK1) | DB04272 DB01992 |
| 78 | succinate dehydrogenase flavoprotein subunit | [Q63JJ8](https://www.uniprot.org/uniprot/Q63JJ8) | DB04657 DB04795 DB09270 DB00139 |
| 79 | aconitate hydratase | [Q63JJ0](https://www.uniprot.org/uniprot/Q63JJ0) | DB06757 |
| 80 | fusion protein, ATP-binding transmembrane ABC transporter and regulatory protein | [Q63I10](https://www.uniprot.org/uniprot/Q63I10) | DB05154 |
